# Benchmarking of Low Coverage Sequencing Workflows for Precision Genotyping in Eggplant

**DOI:** 10.1101/2024.10.24.619843

**Authors:** Virginia Baraja-Fonseca, Andrea Arrones, Santiago Vilanova, Mariola Plazas, Jaime Prohens, Aureliano Bombarely, Pietro Gramazio

## Abstract

Low-coverage whole-genome sequencing (lcWGS) presents a cost-effective solution for genotyping, particularly in applications requiring high marker density and reduced costs. In this study, we evaluated lcWGS for eggplant genotyping using eight founder accessions from the first eggplant MAGIC population (MEGGIC), testing various sequencing coverages and minimum depth of coverage (DP) thresholds with two SNP callers, Freebayes and GATK. Reference SNP panels were used to estimate the percentage of common biallelic SNPs (i.e, true positives, TP) relative to the low coverage datasets (accuracy) and the SNP panels themselves (sensitivity), along with the percentage of TP with the same genotype across the two datasets (genotypic concordance). Sequencing coverages as low as 1X and 2X achieved high accuracy but lacked sufficient sensitivity and genotypic concordance. However, 3X sequencing reached approximately 10% less sensitivity than 5X while maintaining genotypic concordance above 90% at any DP threshold. Freebayes outperformed GATK in terms of sensitivity and genotypic concordance. Therefore, we used this software to conduct a pilot test with some MEGGIC lines from the fifth generation of selfing (S5), comparing their datasets with a gold standard (GS). Sequencing coverages as low as 1X identified a substantial number of TP, with 3X significantly increasing the yield, particularly at moderate DP thresholds. Additionally, at least 30% of the TP were consistently genotyped in all lines when using coverages greater than 2X, regardless of the DP threshold applied. This study highlights the importance of using a GS to reduce false positives and demonstrates that lcWGS, with proper filtering, is a valuable alternative to high-coverage sequencing for eggplant genotyping, with potential applications to other crops.

## 1. Introduction

Plant genomic characterization is a critical step in modern breeding programs, and essential for studying diversity, understanding domestication and recombination events, and identifying candidate regions linked to important agronomic traits (Song et al., 2023). Current methods for high-throughput genotyping in plants primarily involve two approaches: reduced representation sequencing (RRS) and whole-genome sequencing (WGS) (Scheben et al., 2017). RRS, represented by techniques such as restriction-site associated DNA sequencing (RADseq; Baird et al., 2008), genotyping-by-sequencing (GBS; Elshire et al., 2011), RNA-Seq (Marguerat and Bähler, 2010) or amplicon-sequencing (Campbell et al., 2015), while cost-effective, exhibits limitations in capturing a comprehensive set of informative sites (Pereira-Dias et al., 2019; Wang et al., 2020). On the other hand, WGS offers an exhaustive genome-wide view, but its higher costs, driven by the need for extensive sequencing coverage and reliance on a reference genome, makes it less accessible for large-scale studies (Ratnaparkhe et al., 2020; Ren et al., 2021). To address the limitations of both approaches, low-coverage whole-genome sequencing (lcWGS) is gaining popularity as a genotyping strategy that combines the broad genomic coverage of WGS with the cost-efficiency of RRS (Golicz et al., 2015; Kumar et al., 2021). lcWGS allows for the identification of a dense marker set with a comprehensive representation of the entire genome, using very low sequencing coverage (<10X) (Malmberg et al., 2018; Happ et al., 2019). This strategy has opened new possibilities for genotyping studies in various crops, including chickpea (Bayer et al., 2015), rice (Wang et al., 2016), canola (Malmberg et al., 2018), soybean (Happ et al., 2019), tomato (Gonda et al., 2019), radish (Luo et al., 2020), wheat (Adhikari et al., 2022) and potato (Clot et al., 2024) among others.

The primary strength of lcWGS lies in its cost-effectiveness, with costs scaling down in tandem with reduced sequencing coverage (Kumar et al., 2021; Lou et al., 2021). Another valuable feature of this strategy is the concurrent reduction in data volume as sequencing coverage diminishes, expediting and streamlining bioinformatic analysis (Malmberg et al., 2018; Deng et al., 2022). Nevertheless, low sequencing coverages can lead to erroneous conclusions due to the limited information provided by the low number of reads. Some potential challenges encompass: (1) genotype misclassification, (2) loss of genuine polymorphism, and (3) sequencing errors being erroneously classified as genetic variants (Meisner and Albrechtsen, 2018). To address these drawbacks, rigorous SNP filtering steps are critical (O’Leary et al., 2018; Lou et al., 2021). Furthermore, employing additional procedures is advised for the elimination of false positive calls, such as using more than one SNP calling software (Wickland et al., 2017; Yao et al., 2020) and validating polymorphisms against a set of truly-assumed genetic variants (gold standard; GS), supported by a higher number of reads or validated in various independent studies (Hardwick et al., 2017; Poplin et al., 2017; Zook et al., 2020).

Eggplant (*Solanum melongena* L.) is an economically significant crop ranking as the third most important solanaceous crop after potato and tomato in global production and the fifth among all vegetable crops (FAOSTAT, 2024). Despite its economic importance, available genetic and genomic resources of eggplant have traditionally lagged behind those of other important vegetable crops (Gramazio et al., 2023). These limitations have hindered genetic advancements, including the identification of quantitative trait loci (QTLs) and causative genes for traits of interest in breeding, the development and exploitation of experimental populations, and the performing of resequencing studies. However, noteworthy progress includes the availability of high-quality reference genomes for eggplant (Barchi et al., 2019b, 2021; Wei et al., 2020; Li et al., 2021), along with the development of biparental populations, such as sets of introgression lines (ILs) (Mennella et al., 2010; Gramazio et al., 2017; Plazas et al., 2020), and the first and the only multiparent advanced generation inter-cross (MAGIC) population developed so far in eggplant, known as MEGGIC (Mangino et al., 2022).

The use of multiple founders in MAGIC populations widens the genetic and phenotypic base of the crop, increasing the number of segregating QTLs due to the large number of accumulated recombinant events that improve QTL mapping accuracy (Arrones et al., 2020). Nevertheless, to fully leverage MAGIC populations as powerful next-generation genomic resources, genotypic characterization is essential. For the MEGGIC population, the 5k SPET (Single Primer Enrichment Technology) genotyping platform (Barchi et al., 2019a), developed from the resequencing at 20X of its eight founders (Gramazio et al., 2019), was used to genotype 420 individuals from the third generation of selfing (S3MEGGIC), resulting in 7,724 high-confidence SNPs (Mangino et al., 2022). Even though the SPET genotyping allowed the dissection of key genes for eggplant genetics and breeding (Arrones et al., 2022, 2024; Mangino et al., 2022), the genetic characterization of the segregating individuals through variant identification and haplotype resolution was not fully comprehensive. Two principal limitations of this genotyping approach by amplicon sequencing are the limited number of variants that can be interrogated and their distribution, which are preferentially selected in the gene-rich chromosome arms to assess gene allelic diversity (Barchi et al., 2019a). These issues can be addressed by lcWGS, as demonstrated in recent specific eggplant studies that created high-density recombination bin-based genetic maps and improved QTL mapping resolution (Qian et al., 2021; Guan et al., 2022). However, there remains a limited understanding of the impact of several parameters during data processing on its accuracy and sensitivity.

Thus, the principal aim of this study is to establish an optimized workflow for analysing low-coverage genomic data in eggplant using the MEGGIC founders by benchmarking different combinations of sequencing coverages, minimum depth of coverage (DP) filters and SNP callers. A proof-of-concept validation of these findings was conducted on recombinant S5 MAGIC lines of the eggplant MAGIC population (S5MEGGIC), which will facilitate the optimization of genomic diversity analyses in eggplant collections and populations. Furthermore, this study provides guidelines for selecting appropriate parameters in eggplant genomics analysis and presents a protocol that can be broadly applied across various crops and research objectives.

## 2. Materials and methods

### 2.1. Plant materials, library preparation and resequencing

The plant materials used for the low-coverage sequencing benchmarking were the eight founders of the MEGGIC population, consisting of seven *Solanum melongena* and one wild relative *S. incanum* accessions (Supplementary Data S1; Mangino et al., 2022). The whole genomes of these founders were previously resequenced at around 20X to capture their genetic diversity (Gramazio et al., 2019). To increase the genomic characterization precision of the MEGGIC population and to validate the lcWGS benchmarking results of this study, four random recombinant lines from the S5 generation (labelled for this study as S5-1, S5-2, S5-3, S5-4) were used. The S5 lines were obtained following a funnel scheme as described by Mangino et al. (2022) (Supplementary Data S1).

Seeds from the twelve samples (the eight founders and the four S5MEGGIC lines) were germinated in Petri dishes, following the protocol developed by Ranil et al. (2015). They were subsequently transferred to seedling trays in a climatic chamber under a photoperiod and temperature regime of 16 h light (25 °C, 100–112 μmol m−2 s−1) and 8 h dark (18 °C). Total genomic DNA was extracted from approximately 100 mg of young leaves following the SILEX protocol described by Vilanova et al. (2020). DNA integrity and quality of the extracted DNA were assessed through agarose electrophoresis and NanoDrop ND-1000 spectrophotometer (NanoDrop Technologies, Wilmington, Delaware, USA). DNA concentration was determined with a Qubit® 2.0 fluorometer (Thermo Fisher Scientific, Waltham, MA, USA). High-quality DNA samples (260/280 and 260/230 ratios > 1.8) were then shipped to the Beijing Genomics Institute (BGI Genomics, Hong Kong, China) for the construction of 150 bp paired-end libraries and subsequent sequencing using the DNBseq platform. The lcWGS was pursued to generate around 6.3 Gb high-quality sequence reads per sample, approximating 5X genome coverage. Raw reads underwent filtration using SOAPnuke software (Chen et al., 2018) to remove adapters and low-quality reads (-n 0.001 -l 10 -q 0.4 --adaMR 0.25 --ada_trim). After receiving trimmed reads, quality control was performed using FastQC (version 0.11.9; Andrews, 2010) to assess the effectiveness of the quality filtering (Figure 1.A and Figure 1.B).

**Figure 1.**
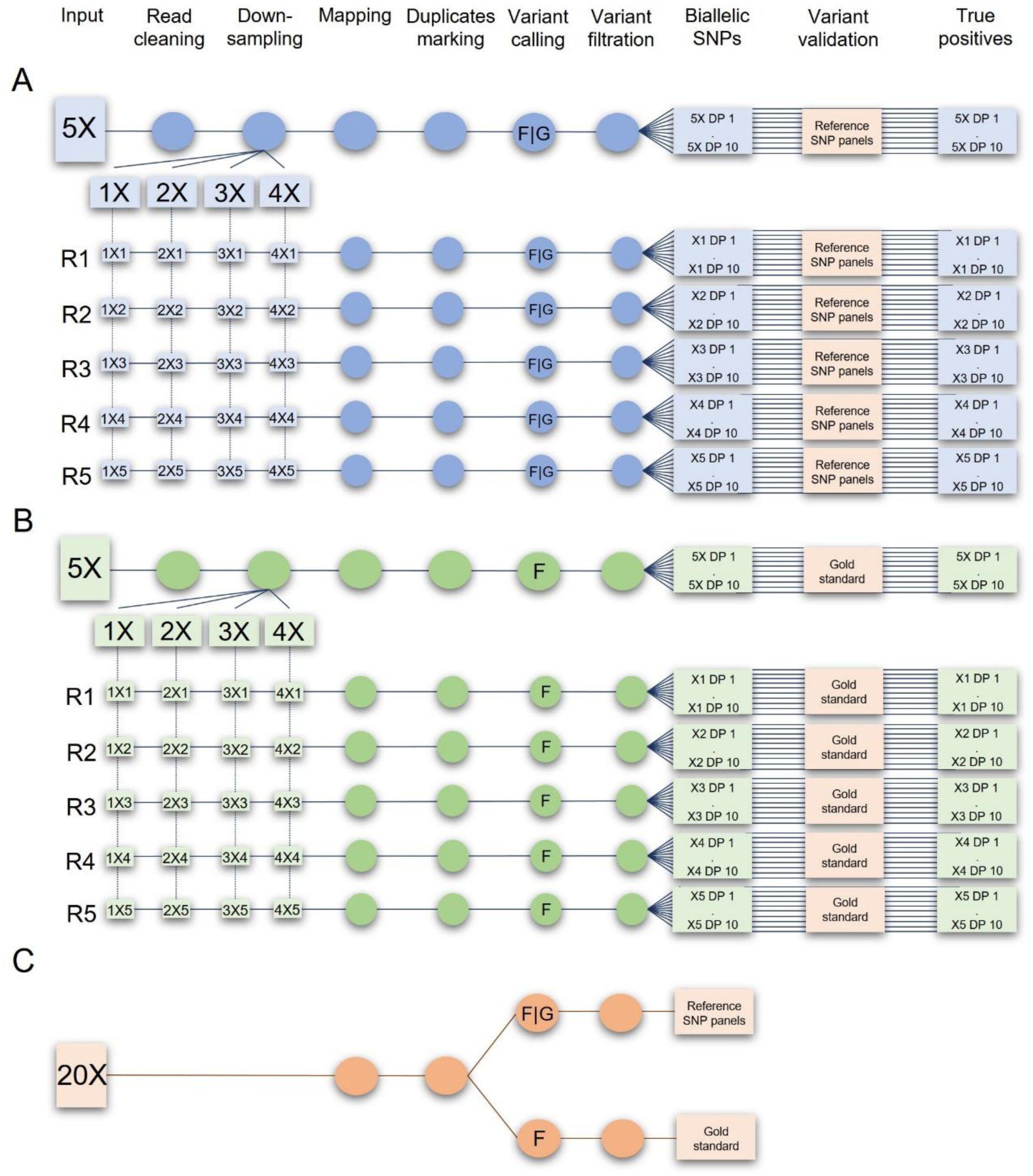
Bioinformatics pipeline for low-coverage genomic data analysis. Squares represent files and circles indicate procedures during the bioinformatic analysis. **A.** Low-coverage whole genome sequencing benchmark workflow. MEGGIC founders’ 5X cleaned data were down-sampled to 1X, 2X, 3X and 4X. Five replicates were generated for each down-sampled level (R1-R5). After the mapping step, SNP calling was performed using Freebayes (F) and GATK (G), followed by variant filtration based on the minimum depth of coverage (DP 1 to DP 10). The SNPs datasets were validated using the corresponding reference SNP panel. **B.** Proof-of-concept validation of the benchmark study. S5MEGGIC lines’ 5X cleaned data were downsampled to 1X-4X, and five replicates were generated for each level (R1-R5). Following the mapping step and the population-level SNP calling with Freebayes, variant filtration was based on the minimum depth of coverage (DP 1 to DP 10). The SNP datasets were validated using the gold standard (GS). Output datasets were labelled according to the sequencing coverage and the applied filter (e.g., 1X DP 1 indicates a sequencing coverage of 1X and a minimum depth of coverage threshold set at 1). **C.** Reference SNP panels and GS preparation from MEGGIC founders’ 20X data. One reference SNP panel per genotype-SNP caller (Freebayes, F; GATK, G) combination was obtained. Additionally, a unique GS was obtained by performing a population-level SNP calling using Freebayes followed by variant filtration.

### 2.2. Downsampling and mapping

The original fastq files obtained from the 5X sequencing were utilized to simulate different lower-depth samples using seqtk tool (version 1.3-r106, https://github.com/lh3/seqtk). Each sample was computationally subsetted based on the number of reads. Average depths of 1X, 2X, 3X and 4X were produced (Figure 1.A and Figure 1.B). The same random seed (-s) was employed to preserve read pairing. For each simulated dataset, five replicates (R) were randomly generated (Figure 1.A and Figure 1.B).

Clean reads from the five gradient sequencing coverages for each sample were mapped against the v3.0 “67/3” high-quality eggplant reference genome (Barchi et al., 2019b) using BWA with its Minimal Exact Match algorithm (BWA-MEM) (version v.0.7.17–r1188; Li, 2013; Figure 1.A and Figure 1.B). The resulting alignment data were subsequently transformed into the BAM format using the SAMtools package (version 1.13; Li et al., 2009). Mapping statistics, including the number of mapped and unmapped reads, and the average depth of coverage, were recorded in the output file generated by the QualiMap application (version 2.2.1; García-Alcalde et al., 2012). Genome coverage was calculated with the ‘coverage’ function within SAMtools package (version 1.13; Danecek et al., 2021). In-depth analysis of the spatial distribution of mapping depth across the genome, with a window size of 10 Kbp, was performed with the bamCoverage tool (version 3.5.1; Ramírez et al., 2014). Output data was visualized and graphically represented using the ‘plot’ function (version 3.6.2) in R (version 4.3.2; (R Core Team, 2021). Finally, PCR duplicates were marked using the MarkDuplicates tool from Picard software (version 1.119; https://broadinstitute.github.io/picard/; Figure 1..A and Figure 1.B).

### 2.3. Polymorphism detection and data filtering

Variant calling in low-coverage founders’ data was carried out at the sample level (i.e., each sample independently for each combination of sequencing coverage and depth) using two different Bayesian-based software: Freebayes (version 1.3.6; Garrison and Marth, 2012) and GATK HaplotypeCaller (version 4.3.0.0; McKenna et al., 2010) (Figure 1.A). Both callers were launched with default configuration settings, except for the minimum quality requirements for mapping and base, which were set at a threshold of 20. Biallelic SNPs, excluding monomorphic ones, were kept using BCFtools (version 1.13; https://samtools.github.io/bcftools/bcftools.html). An additional filtering step was implemented to assess the impact of the DP thresholds on polymorphism detection, ranging from DP 1 to DP 10. This process generated a total of 210 list-based SNP sets per sample-SNP caller combination, capturing variations across different depth levels. Specifically, this includes five replicates for each combination of 10 minimum depth coverages across sequencing coverages from 1X to 4X, along with a single set for each depth at 5X coverage (Figure 1.A).

Variants of the four S5MEGGIC lines were identified at the population level (i.e., the four samples together) using Freebayes (Figure 1.B). While the default configuration settings were primarily maintained, the thresholds for mapping and base quality were again specifically set at a minimum of 20. In the same way, as did with founders’ data, biallelic SNPs were filtered by the minimum depth of coverage ranging from DP 1 to DP 10. Finally, only polymorphic variants among the four lines were retained (Figure 1.B).

### 2.4. MEGGIC founders derived reference SNP panels and gold standard

To benchmark each MEGGIC founder’s combinations of lc-DP (low-coverage dataset and minimum depth of coverage threshold), two reference SNP panels were established for each MEGGIC founder from its 20X resequencing dataset (SRA BioProject PRJNA392603; Gramazio et al., 2019) (Figure 1.C). The cleaned reads underwent a quality control assessment using FastQC (version 0.11.9; Andrews, 2010). The mapping and polymorphism detection steps using Freebayes (Freebayes SNP panel) and GATK (GATK SNP panel) were conducted as previously detailed for the low-coverage founders’ data. Biallelic SNPs supported by at least 20 reads were retained to ensure the most informative markers, while monomorphic ones were excluded (Figure 1.C).

In addition, a GS was established to select common biallelic SNPs identified in the S5MEGGIC lines, thereby validating the benchmark results and determining the optimal combination of parameters for the genomic characterization of the S5MEGGIC population (Figure 1.C). To set up the GS, SNP calling was performed at the population level using the 20X resequencing datasets from all eight founders with Freebayes (-q 20 -m 20 –limit-coverage 800), generating a unique VCF file that contained both common and unique variants across all founder genomes. Biallelic SNPs supported by at least 20 reads were retained, and monomorphic sites among samples were removed (Figure 1.C).

### 2.5. Genotyping accuracy, sensitivity and concordance evaluation

To determine the optimal combination of tools and parameters in terms of variant discovery, for each founder, the Freebayes and GATK SNP panels were compared with their respective low-coverage datasets (lc-datasets) generated by Freebayes and GATK, respectively (Figure 1.A). These comparisons were performed with the isec tool from the BCFtools package (version 1.13; https://samtools.github.io/bcftools/bcftools.html) with the default configuration, yielding only records with identical alleles between the reference SNP panel and the lc-dataset (bcftools isec -c none). The output files allowed the identification of polymorphic sites: (I) common between the reference SNP panel and the lc-dataset (TP), (II) private to the lc-dataset (FP), (III) and private to the reference SNP panel (PP). The isec tool was also employed to accurately determine the common variants between the S5MEGGIC lines and the GS (TP) (Figure 1.B).

The first metric evaluated was accuracy, defined as the capability to filter out correctly potential false polymorphisms or identification errors. It was calculated as the ratio of TP to the total number of polymorphisms identified in the sample (1).

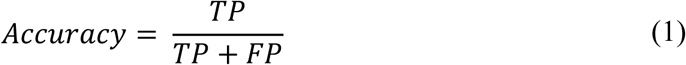

On the other hand, sensitivity was assessed as the capability to identify correctly genuine polymorphisms within the genome. It was calculated as the ratio of TP to the total number of polymorphisms identified in the reference SNP panels (2).

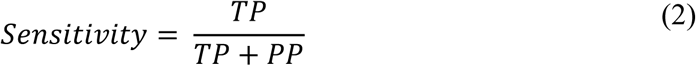

Finally, genotypic concordance was calculated as the percentage of TP in the lc-dataset that matched the genotype at the same site sequenced at 20X coverage. This measure reflected the ability to assign accurately genotypes even at low coverages.

## 3. Results

### 3.1. lcWGS and mapping

The 5X lcWGS of the seven *S. melongena* and one *S. incanum* founders of the MEGGIC population, along with four recombinant S5MEGGIC lines, yielded an average of 19.70 M reads per sample (Supplementary Data S2). Of these, 96.38% were high-quality nucleotide bases, with a Q score greater than 20 (>Q20) (Supplementary Data S2). To benchmark the impact of varying sequencing coverages, subsets ranging from 1X to 4X were simulated (Figure 1.A and Figure 1.B). The original 5X and low-coverage datasets were aligned against the “67/3” eggplant reference genome using BWA-MEM software. The average percentage of mapped reads was 96.40%, with no significant differences observed between the relative data of the 5X and low-coverage sets (Supplementary Data S3). The BAM files derived from the original 5X sequences exhibited an average depth of coverage of 4.79 and the 1X, 2X, 3X, and 4X subsets of 0.96, 1.92, 2.87, and 3.83, respectively (Supplementary Data S3).

As the average sequencing coverage increased, the proportion of the reference genome covered also expanded, although the increments became progressively smaller at higher coverages (Figure 2.A and Supplementary Data S3). At 1X, 39.06% of the reference genome was covered, increasing to 55.20% at 2X and 63.05% at 3X. However, the gains diminished significantly at higher coverages, with only 66.54% and 68.52% covered at 4X and 5X, respectively (Figure 2.A). Surprisingly, a marginal increment of 4.58% was observed from 5X to 20X coverage, covering 71.66% of the reference genome (Figure 2.A).

**Figure 2.**
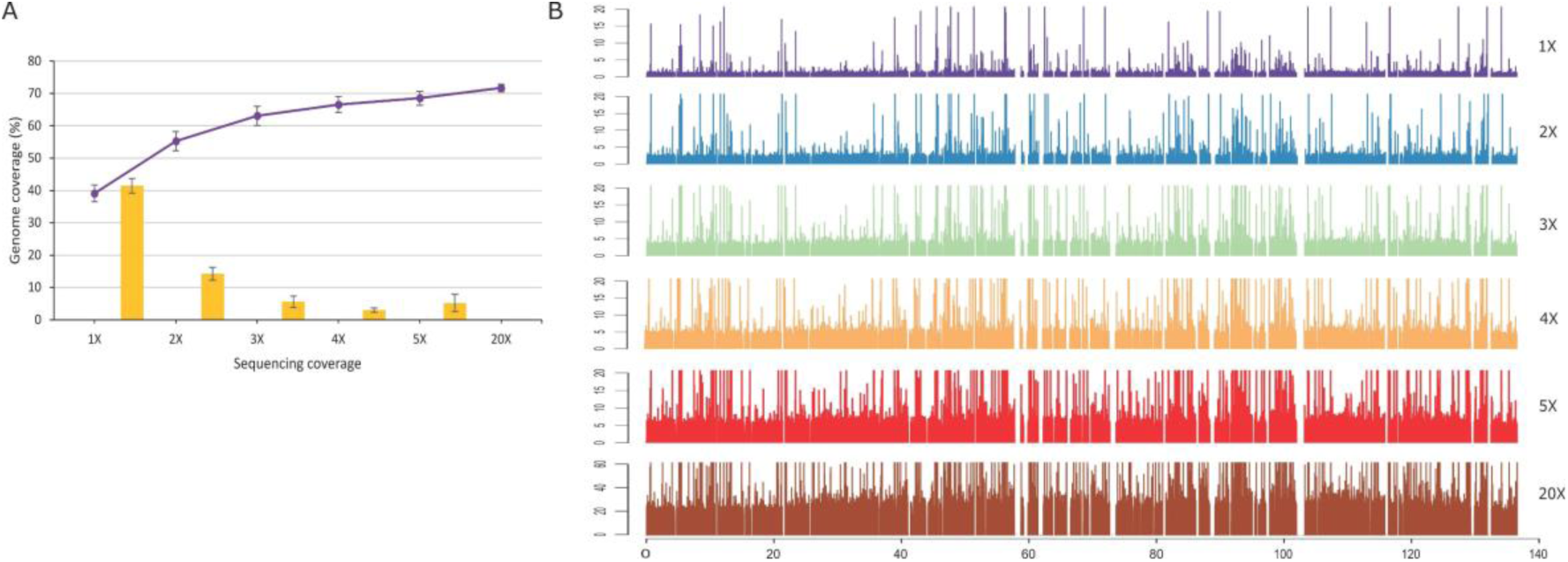
**A.** Percentage of reference genome coverage at different sequencing coverages (1X to 5X, and 20X) and the increments (yellow bars) from one coverage to the next. Genome coverage refers to the percentage of the genome covered by at least one read. The data shown are averages calculated from five replicates of each genotype for 1-4X sequencing coverage and from the original datasets for each genotype for 5X and 20X coverage. Standard deviation is indicated by error bars (n=60 for 1-4X, n=12 for 5X and 20X). **B.** Example of the distribution of mapped read across the parental A chromosome 1 by sequencing coverage. The maximum cutoff is set at DP 20 for low coverages and DP 60 for 20X data. Peaks represent regions of high sequencing within 10 Kbp windows.

The distribution of mapped reads across the sequencing coverages displayed a consistent pattern, characterized by regions of the genome lacking sequencing data (underserved regions) alongside areas subjected to excessive sequencing (over-sequenced regions) (Figure 2.B and Supplementary Data S4).

### 3.2. In silico evaluation of lcWGR under varying parameters

To assess the effectiveness of lcWGS for polymorphism identification, we systematically tested several combinations of sequencing coverages (1X to 5X), depth thresholds (DP 1 to DP 10), and two widely-used SNP callers, Freebayes and GATK. These comparisons aimed to determine which SNP caller and parameters combination provided the best balance between polymorphism yield and resource efficiency.

Regarding SNP caller performance, four key trends emerged from our analysis: (I) Freebayes identified more polymorphisms than GATK, particularly at higher sequencing coverages and lower DP thresholds (Supplementary Data S5 and Supplementary Data S6); (II) while Freebayes showed no significant differences between filtering at DP 1 and DP 2, GATK exhibited a slight reduction in biallelic SNPs, particularly at lower sequencing coverages (Figure 3.A and Supplementary Data S6); (III) the reduction in the number of SNPs identified from DP 1 to DP 10 was more pronounced in GATK (88.92%) compared to Freebayes (81.63%), indicating a more aggressive filtering effect in GATK at higher DP thresholds (Figure 3.A); and (IV) GATK exhibited a higher scaling factor between sequencing coverages (i.e., a more significant proportion of biallelic SNPs identified at each sequencing coverage relative to the previous one, e.g. 2X:1X), identifying a larger proportion of new SNPs as coverage increased from 2X to 3X (2.16) and 4X to 5X (1.45), compared to Freebayes (1.97 and 1.34, respectively) (Figure 3.B). However, Freebayes slightly surpassed GATK when moving from 1X to 2X (3.31 vs 3.29) (Figure 3.B).

**Figure 3.**
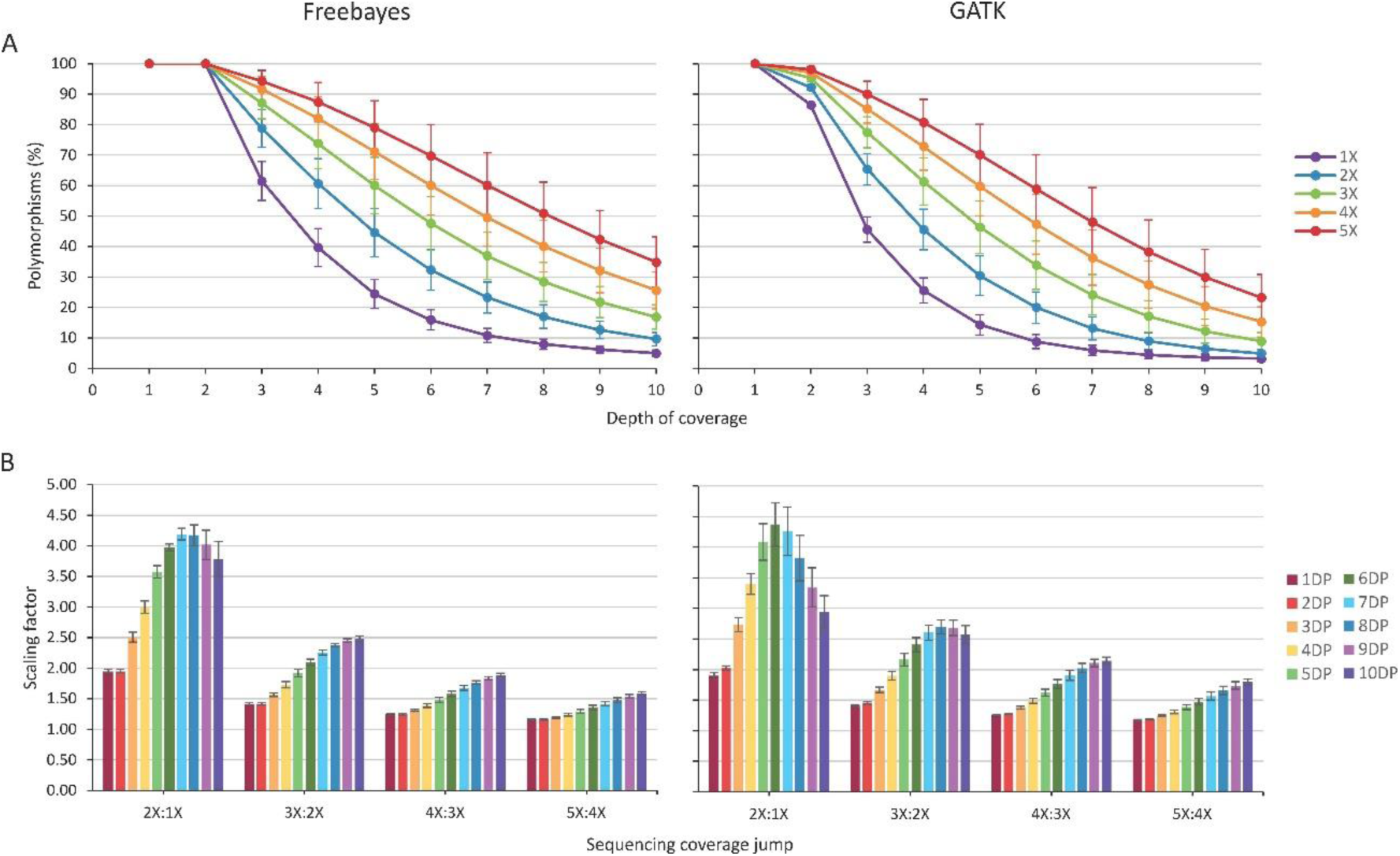
**A.** Variation in the percentage of total biallelic SNPs when using different minimum depth of coverage thresholds (from DP 2 to DP 10) compared to DP 1 (set at 100%) for different sequencing coverages (1-5X) using Freebayes and GATK. Percentages represent the average across the eight founder accessions, with SD indicated by error bars (n=40 for 1-4X and n=8 for 5X). **B.** Variation in the scaling factor, indicating the change in the number of total biallelic SNPs when increasing the sequencing coverage from one coverage to the next, using different minimum depth of coverage thresholds (from DP 1 to DP 10) with Freebayes and GATK. Values represent the average across the eight founder accessions, with SD indicated by error bars (n=40).

Both sequencing coverage and DP threshold had a significant influence on the number of polymorphisms identified, with higher coverage and lower DP thresholds consistently yielding more biallelic SNPs (Supplementary Data S6). However, the impact of DP threshold was dependent on the sequencing coverage. At lower coverage, the SNP reduction was more pronounced at lower DP thresholds than at higher ones (Figure 3.A). For example, at 1X with Freebayes, the SNP reduction from DP 2 to DP 3 was 38.54%, whereas the difference between DP 9 and DP 10 was only 1.15% (Figure 3.A). Conversely, at higher sequencing coverages, the impact of DP thresholds on SNP reduction was less significant. For instance, at 5X with Freebayes, filtering at DP 3 resulted in a 5.71% reduction compared to DP 2, which was closer to the 7.48% difference observed between filtering at DP 9 and DP 10 (Figure 3.A).

This trend was also observed in the scaling factor when transitioning from one sequencing coverage level to the next. When comparing the number of SNPs identified at 1X and 2X (2X:1X), the scaling factor was around 2 at DP 1 and DP 2, indicating that at 2X, the number of SNPs identified was double that at 1X (Figure 3.B). At higher DP thresholds, the scaling factor increased to 3 or 4 with both Freebayes and GATK (Figure 3.B). In contrast, the scaling factor was more consistent across DP thresholds at higher sequencing coverages. For example, when moving from 4X to 5X, the scaling factor ranged from 1.16 at DP 1 to 1.59 at DP 10 using Freebayes, and from 1.18 to 1.80 with GATK (Figure 3.B). Additionally, at lower sequencing coverages, a plateau was achieved for the scaling factor, but this was not observed at higher coverages (5X:4X), where the scaling factor increased steadily across DP thresholds.

### 3.3. Prediction of the optimal combination of factors to perform lcWGS

Comparisons between the reference SNP panels and each SNP dataset across lc-DP combinations allowed for the determination of TP, missing data, and genotype assignment errors, as well as the estimation of accuracy and sensitivity. These panels were generated from the 20X resequencing dataset of the MEGGIC founders (Gramazio et al., 2019), assuming that the genotypes were more accurately characterized due to the higher coverage. The total cohorts of biallelic SNPs constituting the Freebayes SNP panels ranged from 3.68 M to 7.59 M (Supplementary Data S7). On average, these were 1.58 times greater than those identified by GATK, which ranged from 1.86 M to 4.93 M (Supplementary Data S7).

For both SNP callers, we evaluated 210 SNP datasets generated from 5X downsampling, observing a consistent increase in TP with higher sequencing coverage and a reduction with stricter DP thresholds (Supplementary Data S6). Freebayes consistently identified more TP than GATK across all coverage levels, which aligned with results observed at 20X (Supplementary Data S7). For instance, at DP 1, Freebayes outperformed GATK, identifying an additional 231.20 k TP at 1X, 574.71 k at 3X and 3.26 M at 5X. Similarly, at the more stringent DP 10 threshold, Freebayes detected more TP than GATK across coverages, with differences ranging from 11.20 k at 1X to 425.03 k at 5X (Supplementary Data S6).

Accuracy trend differed between the two SNP callers. At low DP thresholds (DP 1 to DP 4), Freebayes achieved higher average accuracy than GATK. While Freebayes exhibited values that ranged from 44.82% at 5X DP 1 to 49.87% at 1X DP 1, the average accuracy obtained with GATK at the same DP threshold showed less variability among sequencing coverages, ranged from 39.41% at 1X to 39.07% at 5X (Figure 4). However, at higher DP thresholds, GATK not only surpassed Freebayes in accuracy but also exhibited increased accuracy variability among sequencing coverages. This trend was evident from DP 6 at 1X, continuing to increase up to DP 10 for 5X coverage. At DP 10, Freebayes’ accuracy varied from 55.68% at 1X to 59.58% at 3X and 57.20% at 5X, compared to GATK’s range of 75.65% at 1X, 66.41% at 3X and 58.77% at 5X (Figure 4). Notably, Freebayes achieved its highest values (>59%) at specific DP thresholds: DP 5-6 at 1X, DP 6-9 at 2X and DP 9-10 at 3X. In contrast, GATK did not reach a plateau, as its accuracy continued to increase at higher thresholds, although the rate of improvement diminished as threshold rose (Figure 4). Overall, the peak accuracy for Freebayes was observed at 3X DP 10, while GATK’s best performance was at 1X DP 10. On the contrary, the observed trend in average sensitivity was consistent for both SNP callers, showing a decrease at higher DP thresholds and lower sequencing coverage (Figure 4). In terms of performance, Freebayes outperformed GATK in sensitivity. At 1X, Freebayes average sensitivity varied from 9.88% at DP 1 to 2.85% at DP 5 and 0.54% at DP 10, whereas GATK sensitivity ranged from 8.50% at DP 1 to 1.83% at DP 5 and 0.51% at DP 10. At 5X coverage, Freebayes achieved an even better performance than GATK with 2.30% higher sensitivity values at DP 1 (35.27% vs 32.97% with GATK), 4.30% at DP 5 (30.45% vs 26.16%) and 4.22% at DP 10 (15.36% vs 11.14%) (Figure 4).

**Figure 4.**
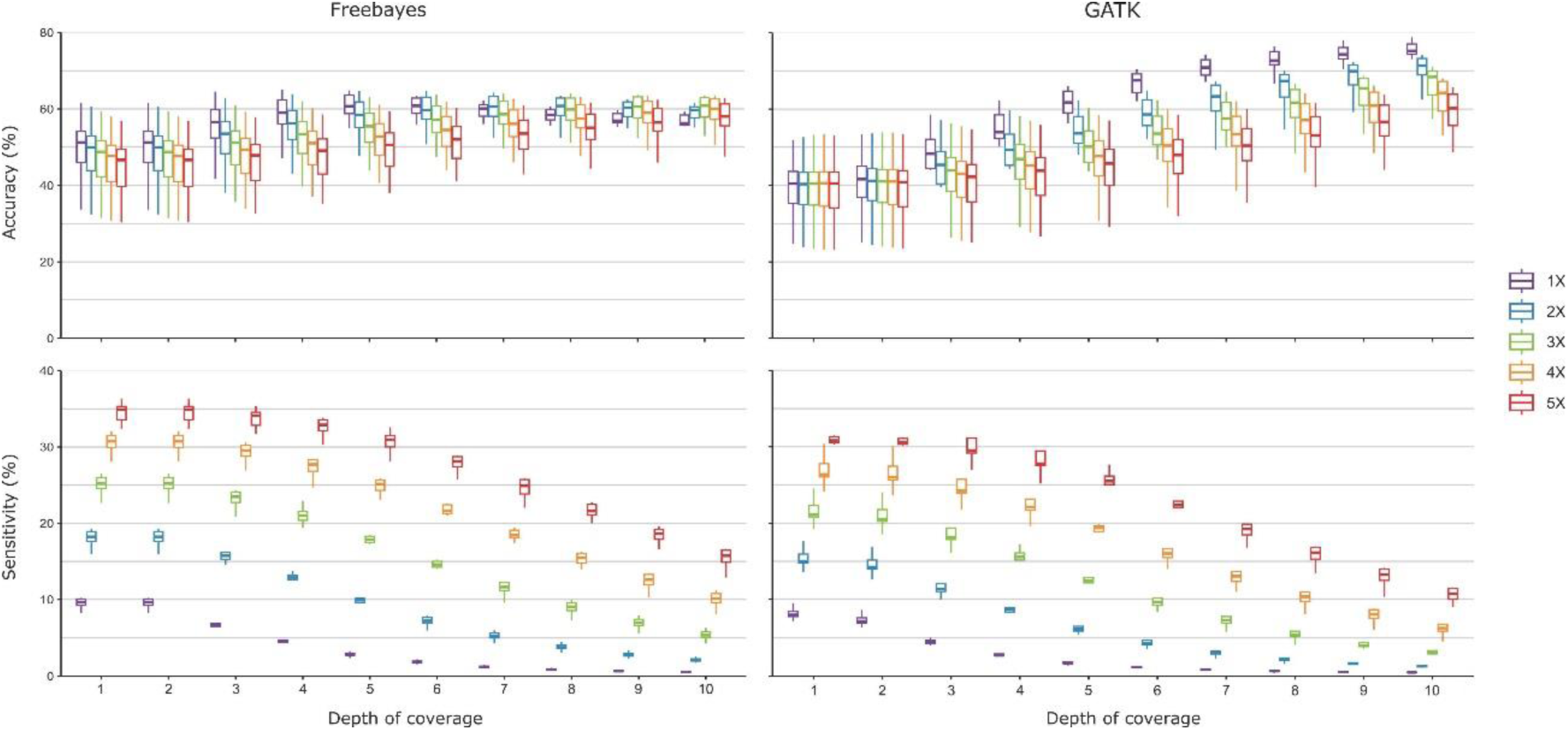
Accuracy and sensitivity (%) achieved for each combination of sequencing coverage (1-5X) and mínimum depth of coverage threshold (from DP 1 to DP 10) using Freebayes and GATK. Accuracy refers to the ability to correctly filter out potential false positives or identification errors, calculated as the ratio of TP to the total number of identified polymorphisms. Sensitivity represents the ability to detect genuine polymorphisms within the genome, calculated as the ratio of TP to the total number of polymorphisms identified in reference SNP panels. A total of 40 samples were evaluated for 1-4X, while 8 samples were assessed for 5X.

Among the common variants identified between the lcWGS datasets and the corresponding reference SNP panels, genotypic concordance was influenced by both SNP caller and sequencing parameters. Freebayes consistently showed higher genotypic concordance under low coverage conditions compared to GATK (Figure 5). For instance, Freebayes identified 65.88% of total TP at 1X DP 1, peaking at over 93.86% at DP 7, while GATK lagged behind, reaching a maximum concordance of around 85.90% at DP 6. At higher coverages (5X), Freebayes maintained superior genotyping concordance, achieving a maximum of 98.72% at DP 10, while GATK’s concordance reached 1.45% less than Freebayes (Figure 5.A).

**Figure 5.**
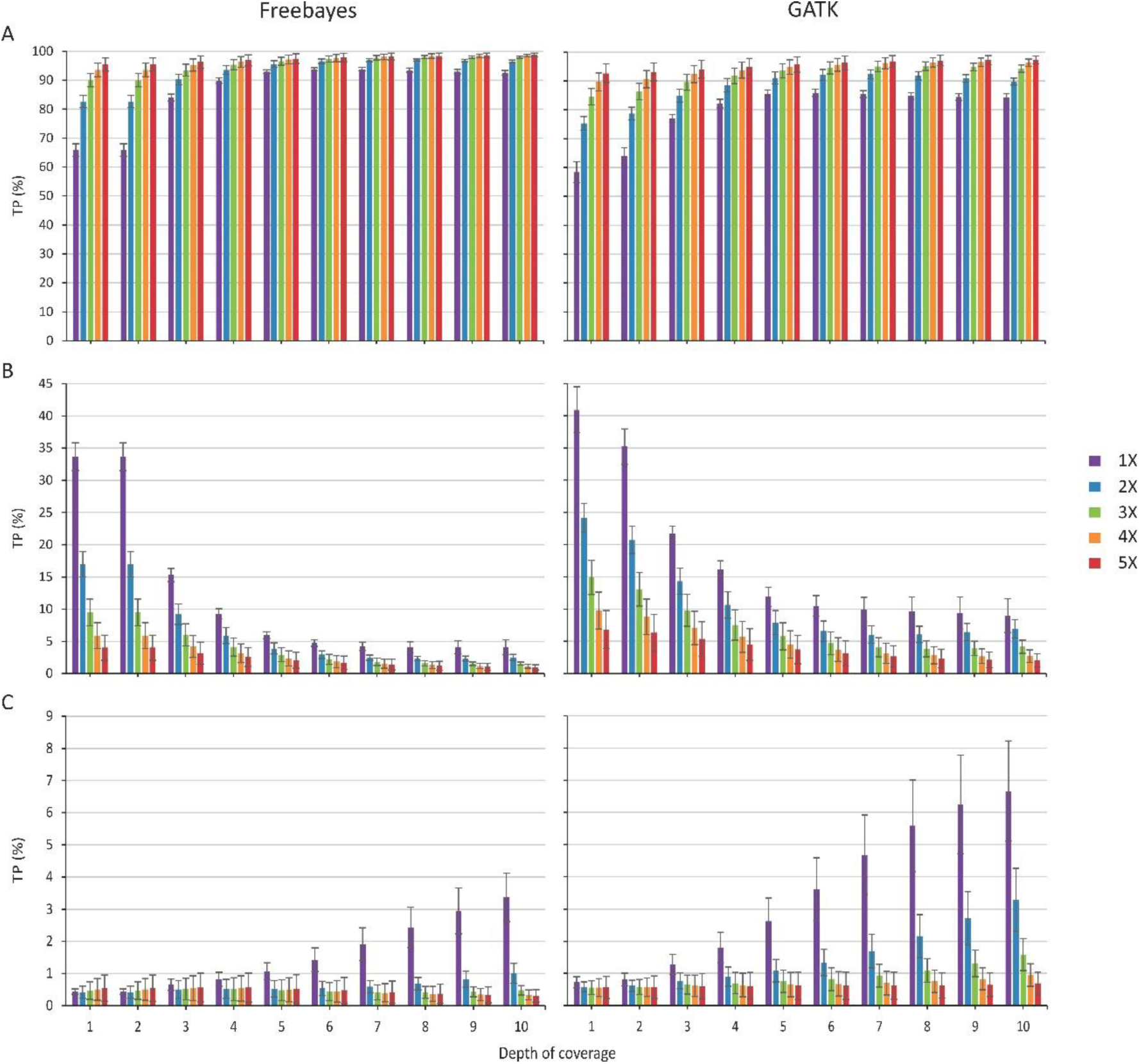
Genotypic concordance and discordance achieved for each combination of sequencing coverage (1-5X) and minimum depth of coverage threshold (from DP 1 to DP 10) using Freebayes and GATK. Percentages represent the average across the eight founder accessions, with SD indicated by error bars (n=40 for 1-4X and n=8 for 5X). **A.** Identical genotypes between the lc-datasets and the reference SNP panels. **B.** Heterozygous sites misclassified as homozygous in the lc-datasets. **C.** Homozygous sites misclassified as heterozygous in the lc-datasets.

Discrepancies between TP and the reference SNP panels genotypes were primarily due to heterozygous loci being misclassified as homozygous in the lc-dataset (Figure 5.B). This error decreased with increasing DP thresholds for both callers. With Freebayes, the percentage of misclassified heterozygous loci fell below 5% starting at DP 5 across all coverages, whereas GATK required higher coverage (3X and above) to achieve similar levels of concordance (Figure 5.B). Homozygous misclassifications were minimal for both callers (Figure 5.C).

### 3.4. Validation of lcWGS results with MAGIC lines

As a proof-of-concept validation of the benchmark performed comparing the MEGGIC founders resequenced at 20X and the corresponding downsampling from 5X, the assessment was extended to four S5MEGGIC lines (Figure 1.B), characterized by an intricate mosaic genome background from the MEGGIC founders (Supplementary Data S1). Similarly, the four S5 lines were sequenced at 5X and downsampled at 4X to 1X with five replicates (Figure 1.B). For the SNP calling, Freebayes was used, guided by insights obtained from comparing the founders’ data with the Freebayes and GATK SNP panels. To assess the extent of potential FP identified at each lc-DP combination, a comparative analysis was conducted using a GS as a reference (Figure 1.B). The GS comprised a combination of shared and private founder variants from the 20X resequencing dataset, obtained by performing a Freebayes SNP calling at the population level (Figure 1.C). This process yielded a total of 17,069,371 biallelic SNPs, distributed according to Supplementary Data S8.

Shared polymorphisms between S5MEGGIC lc-datasets and the GS (i.e., TP) are the most informative in determining the most convenient lc-DP combination to maximize characterization accuracy and resource optimization. As expected, the number of TP identified increased with the sequencing coverage and decreased at higher DP thresholds (Figure 6 and Supplementary Data S9). Similarly to the founders’ benchmark (Figure 3), TP decrements decelerated at higher sequencing coverage. Fixing DP 1 as 100% of TP for each sequencing coverage, 5X DP 10 still retained 21.67% of the TP versus 8.26% at 3X DP 10 and only 1.59% at 1X DP 10 (Supplementary Data S9). Nevertheless, the proportion of TP relative to the total biallelic SNPs identified for each lc-DP combination (%TP) did not follow the same trend and was different for each sequencing coverage (Supplementary Data S10). The %TP slightly increased at higher sequencing coverage and the plateau shifted at higher DP thresholds when adding coverage. So that, at 1X, the higher %TP was 51.76% observed with DP 3, while it was 61.70% at 5X DP 10 (Supplementary Data S10). Thus, considerations should be given to whether applying higher DP thresholds is advantageous, as while it may reduce false positives and increase confidence in calling heterozygous loci, it could also result in a lower proportion of TP relative to the total biallelic SNPs identified.

**Figure 6.**
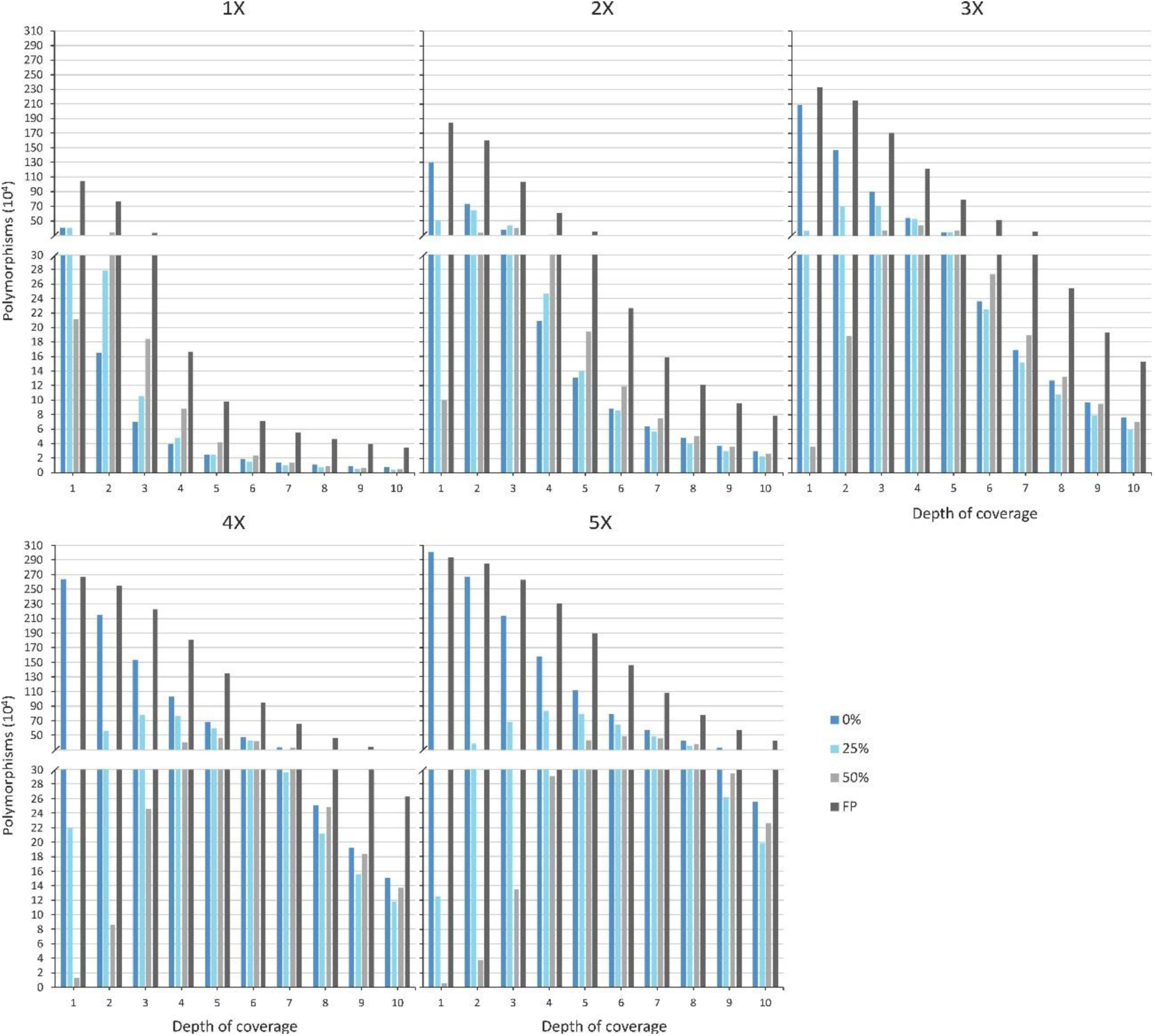
Variation in the total number of polymorphisms when using different minimum depth of coverage thresholds (from DP 1 to DP 10) across different sequencing coverages (1-5X) using Freebayes. The total number of polymorphisms is broken down into false positives (FP) and true positives (TP), with TP being those shared between the samples and the gold standard. TP are categorized based on genotyping completeness in the four S5MEGGIC lines: fully genotyped (0% missing data), partially genotyped in three lines (25% missing data), and in two lines (50% missing data). Values represent the average across the four S5MEGGIC lines, with SD indicated by error bars (n=20 for 1-4X and n=4 for 5X).

Additionally, missing data for each lc-DP combination was assessed, as it is a variable that highly impacts downstream analysis (Figure 6 and Supplementary Data S11). At low DP thresholds (DP ≤ 7), sequencing at 5X achieved the highest percentage of sites without any missing data (95.84% at DP 1). Beyond this threshold, the highest rate was obtained with 1X sequencing coverage. At this coverage, the %TP genotyped in all individuals varied from 39.52% at DP 1 to 45.96% at DP 10, with a minimum value of 19.40% at DP 3, where sites genotyped in two out of four individuals peaked at 51.17% (Supplementary Data S12). As sequencing coverage increased, the minimum percentage of sites genotyped in all individuals moved and was achieved with higher DP thresholds, with a similar trend for the peak of sites genotyped in two individuals. For example, at 3X coverage, the %TP without missing data varied from 83.94% at DP 1 to 37.20% at DP 10, with a minimum value of 32.17% achieved at DP 6. At 5X coverage, it ranged from 95.84% at DP 1 to 37.59% at DP 10 (Supplementary Data S12). Percentage of sites genotyped in three individuals remained more constant across all thresholds, except for high coverages and low DP thresholds (Supplementary Data S12).

## 5. Discussion

Accurately and comprehensively identifying genetic variation is crucial for advancing studies in genetic diversity, trait mapping, and breeding within plant genomics. Experimental populations, such as MAGIC lines, are particularly powerful tools for generating genetic diversity and identifying genomic regions linked to traits of interest (Mangino et al., 2022; Kumar et al., 2023; Thudi et al., 2024). To fully exploit these populations, it is essential to achieve an unbiased and accurate genetic representation of the mosaic genomes present in each line, a goal often constrained by the limitations of the genotyping technology employed (Dell’Acqua et al., 2015; Mangino et al., 2022). In this study, we benchmark the application of lcWGS for high-throughput eggplant genotyping, focusing on both characterizing experimental populations and applying this method to broader genotyping efforts. As lcWGS is relatively novel in plant genomics, we evaluated key parameters —such as sequencing coverage, depth thresholds, and SNP callers —to assess their influence on polymorphism identification. This analysis benefited from the data generated from the WGS of the eight MEGGIC founders, providing a robust foundation for comparison and optimization (Gramazio et al., 2019; Mangino et al., 2022). The results were further validated using S5MEGGIC recombinant fixed lines, offering valuable insights into the broader utility of lcWGS for high-resolution genotyping.

Our results support lcWGS as a promising strategy to address the technical and economic limitations inherent in the main massive genotyping methods used in eggplant, such as SPET (Barchi et al., 2019a), GBS (Peterson et al., 2014), and WGS (Kumawat et al., 2022), offering a synthesis of their respective advantages (Lou et al., 2021). Despite its use of low sequencing coverage, lcWGS provides comprehensive genome representation and identifies a significant number of variants, largely due to the absence of a genome complexity reduction step (Jiang et al., 2019; Wragg et al., 2024). We observed that an increase in the sequencing coverage above 5X resulted in only small increments in genome coverage. Specifically, the difference in reference genome coverage between 5X and 20X was very low. Furthermore, the distribution of mapped reads was similar between the lc-datasets and the 20X data, even though they were generated in different experiments. This implies the presence of regions in the genome that were not sequenced even at high coverage or could not be aligned against the reference genome. This may be due to unassembled regions in the reference genome or the presence of repetitive sequences, which correspond to 12% and 73% of the v3.0 eggplant reference genome “67/3”, respectively (Barchi et al., 2019b).

Several studies have demonstrated that genome coverages as low as 1X are sufficient for association analysis, identifying a significant number of high-quality polymorphisms (Bhattarai et al., 2023; Sapkota et al., 2023; Clot et al., 2024). Even lower sequencing coverages, such as 0.20X and 0.03X, have been successfully used in wheat for precision mapping of key traits (Saripalli et al., 2023), and as low as 0.02X was applied in rice for population characterization and QTL analysis (Huang et al., 2009). In our study, sequencing coverages from 1X to 5X also enabled the identification of a substantial number of SNPs, making it a viable approach for diverse biological analysis, despite the significant percentage of missing loci compared to higher coverage resequencing. As noted in previous work (Malmberg et al., 2018; Adhikari et al., 2022; Deng et al., 2022; Liu et al., 2022), the number of detected SNPs increased with coverage, although the total number varied depending on the applied DP threshold. We explored several DP thresholds (DP 1 to DP 10) as the number of reads supporting a variant directly impacts the reliability of the calls. Raising the DP threshold reduced the number of SNPs, as low-read support variants were filtered out, improving reliability (Malmberg et al., 2018).

Sequencing coverage represents a compromise between cost and the number of polymorphisms detected (Lou et al., 2021; Martin et al., 2021). The additional cost must be weighed against the gain in polymorphisms, which is influenced by the chosen DP threshold. Along with a fixed library preparation cost, the price of sequencing a genome size of ∼3 Gb at 1X coverage is at present around $18, based on Watowich et al. (2023). For the 1.21 GB eggplant genome (Barchi et al., 2021), this translates to around $7 at 1X coverage. Doubling the coverage to 2X doubles the cost, but our results show that this also approximately at least doubled the number of identified polymorphisms across all DP thresholds. Further increases in coverage —such as from 3X to 4X or from 4X to 5X— yielded proportional increases in polymorphism detection starting from a certain DP threshold, but did not double the number of SNPs identified. This aligns with findings by Liu et al., (2022), where the rate of SNP detection slowed as coverage increased. Although increasing coverage does provide more data, this does not necessarily lead to increased accuracy. In fact, we observed similar accuracy across different coverage levels, with a plateau occurring at specific DP thresholds depending on the coverage. This observation is consistent with the work of Song et al., (2016), who reported only a minor 0.5% difference in accuracy between 5X and 10X coverage.

One of the primary challenges in lcWGS is the discrimination between homozygous and heterozygous sites. As noted by Bayer et al., (2015), the correlation between the number of aligned reads and the number of heterozygous SNPs for an individual increased with sequencing coverage. We evaluated genotypic concordance for each lc-DP combination with both SNP callers and observed an improvement as sequencing coverage and DP threshold increased, up to a certain point. This is consistent with findings in animal genomics, observed in species such as *Canis lupus* and *C. familiaris* (Kardos and Waples, 2024; Wragg et al., 2024). With Freebayes, concordance exceeded 90.00% at 3X, 4X and 5X coverage when filtered by any DP threshold. Using GATK, genotypic concordance exceeded 80.00% at these coverages. We found that non-concordant sites were typically misclassified as homozygous when their true genotype was heterozygous, due to the failure to detect both alleles in a small number of sequence reads. Kardos and Waples, (2024) estimated that, at 3X, 4X and 5X coverages, the likelihood of failing to detect one of the two alleles at a heterozygous locus is approximately 0.25, 0.12 and 0.06 in *C. lupus*. Thus, the percentage of heterozygous genotypes at 1X was lower than at 5X, which suggests that heterozygosity was underestimated at low coverages.

When it comes to bioinformatic tools used in lcWGS, the choice of read aligner, while less influential on the accuracy of variant discovery than variant callers, still plays a crucial role in overall data quality (Musich et al., 2021). Among the available options, BWA-MEM has stood out in different benchmarking studies as one of the best read mapper (Wu et al., 2019; Schilbert et al., 2020; Yao et al., 2020). However, the ongoing debate regarding the optimal variant caller in terms of performance remains unresolved (Barbitoff et al., 2022). In this study, we compared two widely used SNP callers, Freebayes and GATK, to determine the best one in terms of the number of TP identified and genotypic concordance when working with lc-data. Freebayes operates as a haplotype-based variant caller detecting polymorphisms based on the sequence content of reads aligned to particular genomic targets (Garrison and Marth, 2012). In contrast, GATK functions as an alignment-based variant caller that detects polymorphisms by locally assembling haplotypes in active regions, relying on precise read alignment to a reference genome (McKenna et al., 2010). In our set of materials, Freebayes identified more TP than GATK at both high and low sequencing coverages, which is in agreement with other studies (Yao et al., 2020; Stegemiller et al., 2023). One potential reason for the superior performance of Freebayes is its utilization of a higher number of reads compared to GATK (Stegemiller et al., 2023). However, this trend is not always consistent. For example, Ni et al., (2015) and Liu et al., (2022) identified 1.31 M and 142.45 k more SNPs using GATK than Freebayes within 8X and 10X whole-genome sequencing data of chicken, respectively.

Regarding the accuracy, sensitivity, and genotypic concordance achieved by each software with various combinations of lc-DP, there were no big differences between the two software. While Freebayes achieved higher sensitivity than GATK, consistent with findings by (Yao et al., 2020), the superiority in accuracy varied depending on the DP threshold used. At higher thresholds, GATK achieved greater accuracy, particularly at low coverages. On the other hand, our results align with Stegemiller et al., (2023), showing that Freebayes assigned more correct genotypes compared to the 20X data than GATK. However, accuracy and sensitivity are also dependent on the specific data used. For example, Ni et al., (2015) found that the set of SNPs obtained from whole-genome sequence data in chicken achieved higher accuracy and genotypic concordance using GATK compared to Freebayes. Similarly, comparisons of variant calling tools for the analysis of *Arabidopsis thaliana* NGS data (Schilbert et al., 2020) and microbial genomes (Bhadhadhara et al., 2023) showed that GATK demonstrated greater accuracy and sensitivity than the other programs evaluated. This is consistent with the study carried out by Liu et al., (2022), which found GATK to be superior at both low and high coverages. In summary, the selection of SNP calling programs should be assessed individually for each specific case. Based on our results, we chose Freebayes for SNP calling the S5MEGGIC lines. We prioritized higher sensitivity and genotypic concordance, as well as time efficiency (Bu et al., 2023), due to the nature of the analysis we conducted.

We also evaluated the use of lcWGS to genomic characterize the S5MEGGIC lines. Due to the additional challenges associated with lc-data, following the same pipeline as with high-coverage data is not advisable. Although the DP threshold allowed us to balance retaining TP with complete genotypic concordance and discarding potential false positives, it may not be sufficient. In the clinical sector, the use of a GS to filter lc-data is widely employed and benefits from a plethora of benchmark datasets (The 1000 Genomes Project Consortium, 2010; Espejo Valle-Inclan et al., 2022). Conversely, its adoption in the breeding sector has been gradual, with reference standards mainly accessible for model species like rice (The 3.000 rice genomes project, 2014), *Arabidopsis* (1001 Genomes Consortium, 2016), corn (Bukowski et al., 2018), soybean (Torkamaneh et al., 2021), and wheat (Jordan et al., 2022). Given the absence of an established GS for eggplant, we developed our own taking advantage of the available 20X data (Gramazio et al., 2019). Unlike SNP calling for the development of reference SNP panels, variant identification within the founders for the GS preparation was conducted in a single step, utilizing all the information collectively. This approach benefited our data as Freebayes leverages information from multiple samples to confidently call variants and address gaps where data from a single sample may be insufficient or ambiguous. More importantly, this method allowed us to obtain complete genotype information for all the samples evaluated from any polymorphic site (Garrison and Marth, 2012). The final dataset used to identify TP in the four S5MEGGIC lines contained 17.07 M SNPs, providing a substantial amount of high-quality data to serve as the GS.

The comparison between the GS and the four S5MEGGIC lines revealed a significant presence of potential FP at low-coverage levels, highlighting the necessity of employing a GS to retain only TP for subsequent analysis (Adhikari et al., 2022; Bhattarai et al., 2023). While the accuracy values, or %TP, were similar between the lc-data from the founders and the lines, the reduction at higher DP thresholds was more pronunced in the lines’ data. This highlights the benefits of considering all lines collectively during SNP calling for the accurate identification of both TP and FP. On the other hand, the application of a DP threshold helped to remove potential FP, but, beyond a certain threshold, TP were also lost. For 1X coverage, a threshold of DP 3 or lower is recommended to capture a useful number of SNPs, while for 3X coverage, filters of less than DP 7 are efficient. Among the TP retained, some genotypes were consistently present across all lines, while others were only genotyped in some, indicating that lcWGS can result in substantial missing genotypes (Malmberg et al., 2018; Kardos and Waples, 2024). Evaluating missing data is essential, as it greatly impacts downstream analyses. For example, tools like the R/mpMap package (Huang and George, 2011) and the magicMap algorithm (Zheng et al., 2019) eliminate markers with excessive missing founder genotypes, given that the number of founder haplotypes increases exponentially with missing data. Although imputation can help address missing genotypes, its accuracy is directly influenced by the proportion of missing data, with higher accuracy achieved as missing data decreases (Nazzicari et al., 2016; Elbasyoni et al., 2018). Our results demonstrated that the percentage of TP with missing data increased at high DP thresholds at 5X coverage. At 1X to 4X coverage, however, this effect was more pronounced at intermediate DP thresholds, likely due to the removal of non-informative sites when applying higher ones. A similar trend may be expected at 5X coverage with more restrictive thresholds.

## 6. Conclusions

Genotyping by lcWGS presents a promising alternative to overcome the limitations of low marker density in GBS and the high costs associated with WGS. While lcWGS is challenged by the identification of numerous potential false positives, our study highlights the efficacy of using a gold standard to mitigate this drawback. Additionally, the choice of bioinformatic tools can significantly influence the results. Future research should focus on optimizing these methodologies and developing tailored bioinformatic tools to fully leverage the potential of lcWGS in various genomic studies. Our findings indicate that Freebayes outperforms GATK in terms of sensitivity and genotypic concordance, though this is data-dependent. Selecting an appropriate sequencing coverage requires careful consideration of multiple variables, including economic constraints, study goals, and species characteristics. We have demonstrated that coverages as low as 1X can yield a substantial number of TP, though higher coverages and stringent threshold depths are recommended for highly heterozygous species to avoid underrepresentation of heterozygosity. Balancing cost, the number of TP, missing data, and genotypic concordance, sequencing coverages of 3X or 4X with threshold depths below DP 7 are optimal. These settings achieve genotypic concordance comparable to 5X while minimizing the loss of TP and the proportion of missing genotypes. Furthermore, combining lcWGS with imputation can effectively address missing genotypes and enhance analytical accuracy, although high levels of missing data can complicate the imputation process. In conclusion, with appropriate optimization, lcWGS presents a cost-effective and scalable tool for genotyping. Successful implementation of lcWGS has the potential to revolutionize genotyping practices not only in eggplant breeding programs but also across other crops and genomic studies, enabling the efficient identification of valuable genetic variants at a reduced cost and accelerating genetic improvement endeavours.

## Acknowledgments

This work was supported by grant PID2021-128148OB-I00 funded by MICIU/AEI/10.13039/501100011033/ and by ERDF/EU, grant CIPROM/2021/020 from Conselleria d’Educació, Cultura, Universitats i Ocupació (Generalitat Valenciana), and by the Horizon Europe programme, project number 101094738 (“Promoting a Plant Genetic Resource Community for Europe; PRO-GRACE). Pietro Gramazio is grateful for the post-doctoral grant RYC2021-031999-I funded by MICIU/AEI/10.13039/ 501100011033 and the European Union through NextGenerationEU/PRTR.

## Data availability

The raw data have been submitted to the NCBI Short Read Archive under the Bioproject identifier PRJNA1174391. Accessions are indexed with BioSample IDs from SAMN44339997 to SAMN44340008. VCF files with the corresponding variants identified are available upon request to the corresponding author.

## Supplementary materials

**Supplementary Data S1. A.** Founders of the MEGGIC population including their country of origin and code assessed in this study. **B.** Funnel breeding design for developing the S5MEGGIC population. The process began with eight parent lines (G0), labelled A to H. These parents were crossed in pairs (A x B, C x D, E x F, G x H) to produce four simple hybrids (G1). Next, these simple hybrids were crossed in pairs (AB x CD, EF x GH) to create two double hybrids (G2). The double hybrids were then crossed (ABCD x EFGH) to form the quadruple hybrids (G3). Following this, the quadruple hybrids underwent intercrossing and were subjected to five rounds of single seed descend (SSD). Adapted from Mangino et al., 2022.

**Supplementary Data S2.** Statistics of the lcWGS of the seven Solanum melongena and one S. incanum founders of the MEGGIC population and for the four S5MEGGIC lines.

**Supplementary Data S3.** Mapping statistics for the individuals assessed in this study. Mean values across five replicates for each level of skim coverage (1-4X) ± SD (n=5).

**Supplementary Data S4**. Distribution of mapped read across the founder accession A by sequencing coverage. The maximum cutoff is set at DP 20 for low coverages and DP 60 for 20X data. Peaks represent regions of high sequencing within 10 Kbp windows.

**Supplementary Data S5**. Comparative heatmaps depicting the average number of biallelic SNPs identified among the MEGGIC founders across varying sequencing coverages (1X to 5X) and minimum depth of coverage thresholds (from DP 1 to DP 10) using Freebayes and GATK. The colour intensity of the squares reflects the number of polymorphisms identified, with darker shades indicating a higher number of biallelic SNPs.

**Supplementary Data S6.** Total number of polymorphic biallelic SNPs for each founder downsampling subsets (1-5X) and minimum depth of coverage thresholds (from DP 1 to DP 10) and shared polymorphisms (TP) with the reference SNP panels from the same founder using Freebayes and GATK. Data from 1X to 4X is the mean value of the five replicates with the standard deviation (SD) reported in the columns to the right.

**Supplementary Data S7.** Reference SNP panels generated by Freebayes and GATK from the 20X resequencing data (Gramazio et al., 2019). “Unfiltered” refers to the biallelic SNPs identified by each SNP caller without applying any filter. Polymorphic biallelic SNPs supported by at least 20 reads constituted the reference SNP panels. The ratio was calculated to compare Freebayes results relative to those of GATK-HC in both cases. Percentages reflect filtered versus total polymorphisms.

**Supplementary Data S8.** Number of missing, referene and alternative genotypes comprising the gold standard for each founder accession. Percentages are calculated above the total 17,069,371 biallelic SNPs.

**Supplementary Data S9.** Total TP identified in the four S5MEGGIC lines and variation in the percentage of total TP (set at 100% at DP 1) across different low sequencing coverage levels (1-5X) and minimum depth of coverage thresholds (from DP 1 to DP 10) using Freebayes. Data for sequencing coverages from 1X to 4X represents the mean value of five replicates. Standard deviations (SD) are shown in the second column.

**Supplementary Data S10.** Percentage of true positives variants when using different minimum depth of coverage thresholds (from DP 1 to DP 10) across different sequencing coverages (1-5X) using Freebayes. True positive polymorphisms were those shared between the samples and the gold standard. Percentages represent the average across the four S5MEGGIC lines, with SD indicated by error bars (n=20 for 1-4X and n=4 for 5X).

**Supplementary Data S11.** Total TP identified in the four S5MEGGIC lines across different percentages of allowed missing data (0%, 25% and 50%), skim sequencing coverage levels (1-5X) and minimum depth of coverage thresholds (from DP 1 to DP 10) using Freebayes. Data for sequencing coverages from 1X to 4X represents the mean value of five replicates. Standard deviations (SD) are shown in the second column.

**Supplementary Data S12.** Percentage of true positives variants identified in the four S5MEGGIC lines (0% missing data), in three lines (25% missing data), and in two lines (50% missing data). This was evaluated for each combination of minimum depth of coverage thresholds (from DP 1 to DP 10) and sequencing coverages (1-5X).

