## Supplementary Data S1 for "Benchmarking of Low Coverage Sequencing Workflows for Precision Genotyping in Eggplant"

**Supplementary Data S1. A.** Founders of the MEGGIC population including their country of origin and code assessed in this study. **B.** Funnel breeding design for developing the S5MEGGIC population. The process began with eight parent lines ($G_0$), labelled A to H. These parents were crossed in pairs (A x B, C x D, E x F, G x H) to produce four simple hybrids ($G_1$). Next, these simple hybrids were crossed in pairs (AB x CD, EF x GH) to create two double hybrids ($G_2$). The double hybrids were then crossed (ABCD x EFGH) to form the quadruple hybrids ($G_3$). Following this, the quadruple hybrids underwent intercrossing and were subjected to five rounds of single seed descend (SSD). Adapted from Mangino et al., 2022.

A

| Taxon and accession | Country of origin | MAGIC's parent code |
|---|---|---|
| *S. incanum* L. | | |
| MM577 | Israel | C |
| *S. melongena* L. | | |
| MM1597 | India | A |
| DH_ECAVI | Breeding line | B |
| AN-S-26 | Spain | D |
| H15 | Spain | E |
| A0416 | Unknown | F |
| IVIA-371 | Spain | G |
| ASI-S-1 | China | H |

B

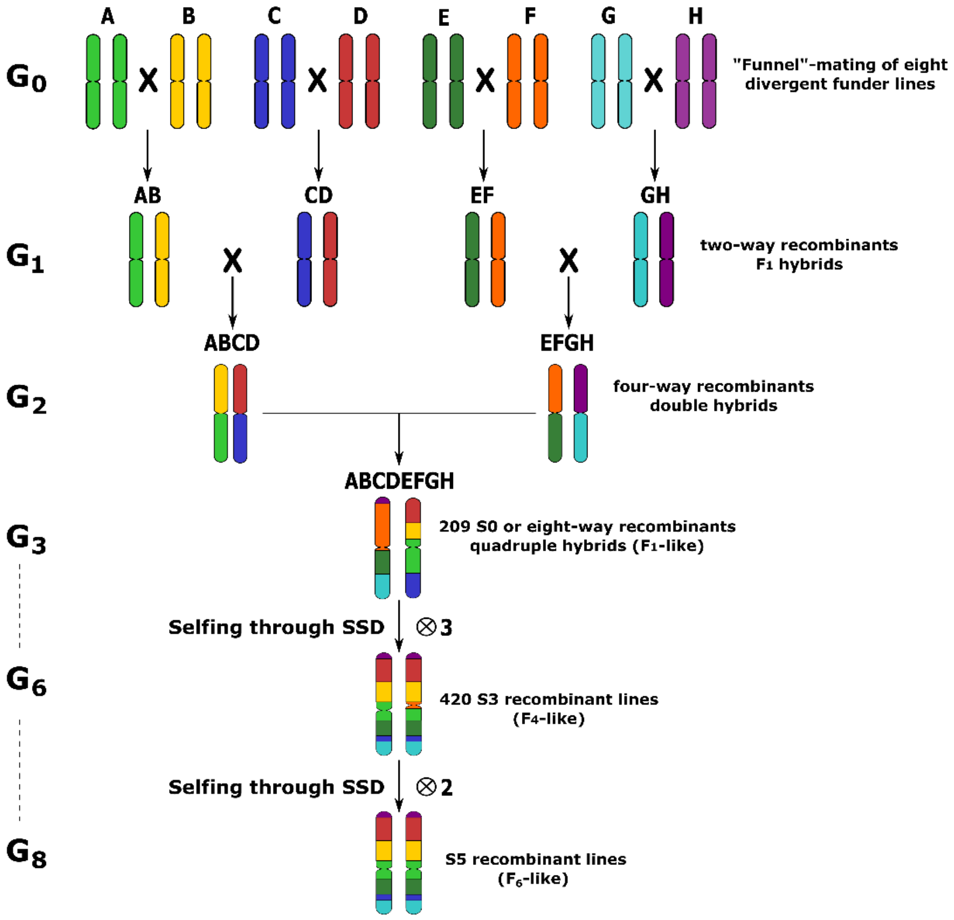
