## Supplementary Data S4 for "Benchmarking of Low Coverage Sequencing Workflows for Precision Genotyping in Eggplant"

**Supplementary Data S4.** Distribution of mapped read across the founder accession A by sequencing coverage. The maximum cutoff is set at DP 20 for low coverages and DP 60 for 20X data. Peaks represent regions of high sequencing within 10 Kbp windows.

Chromosome 2

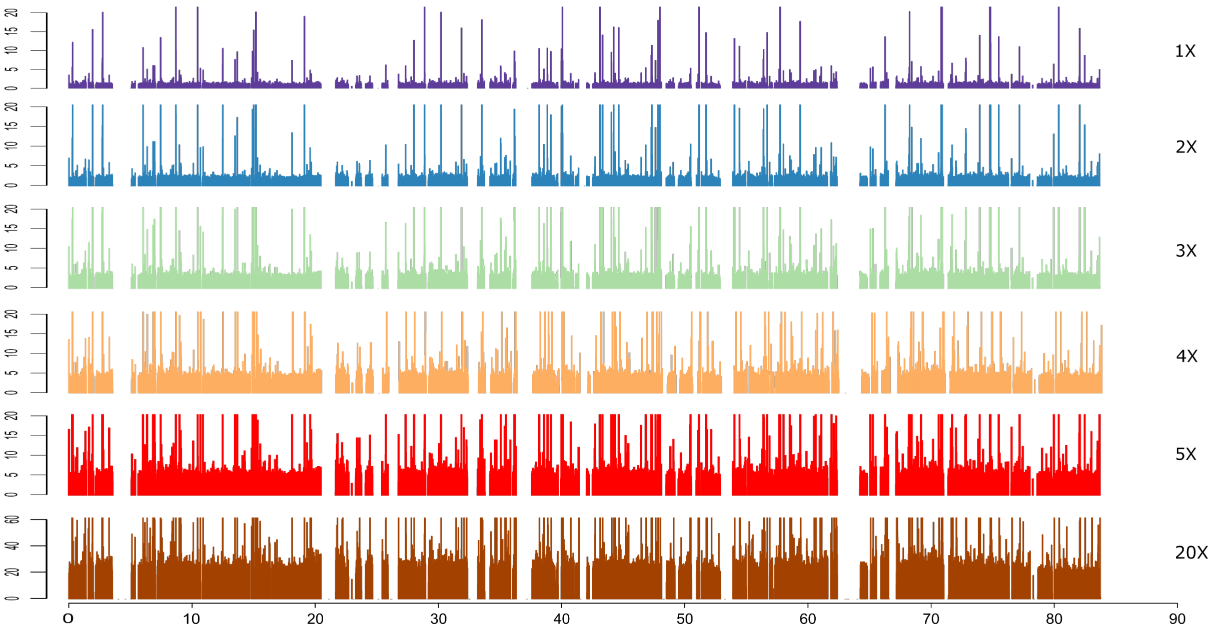


Chromosome 3

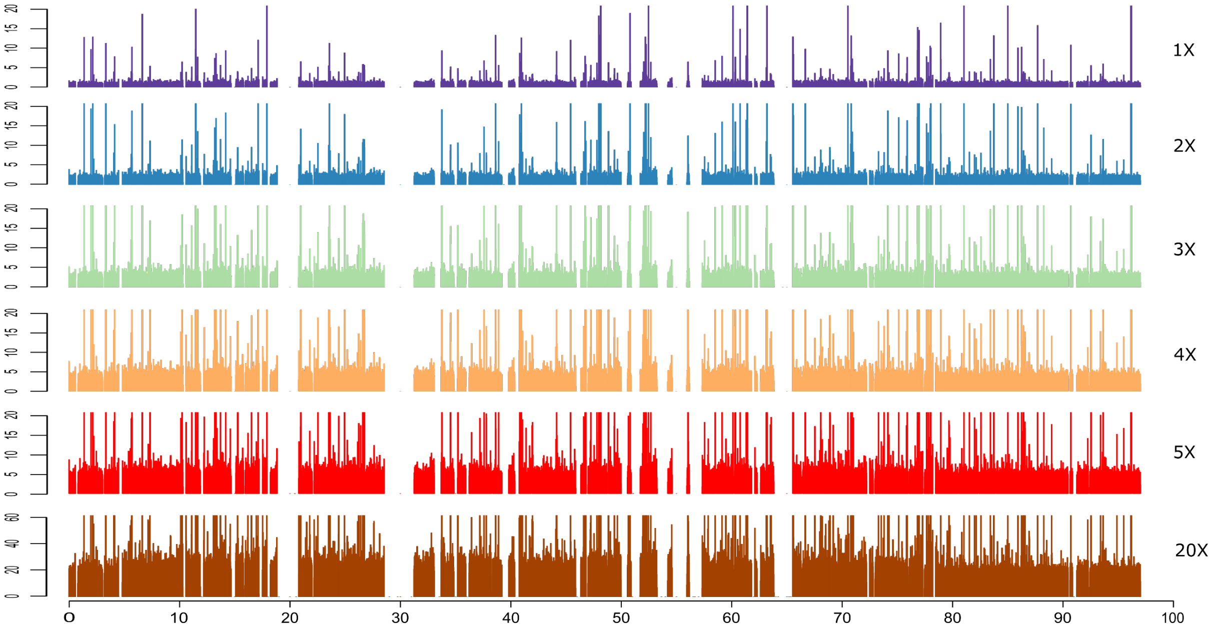

Chromosome 4

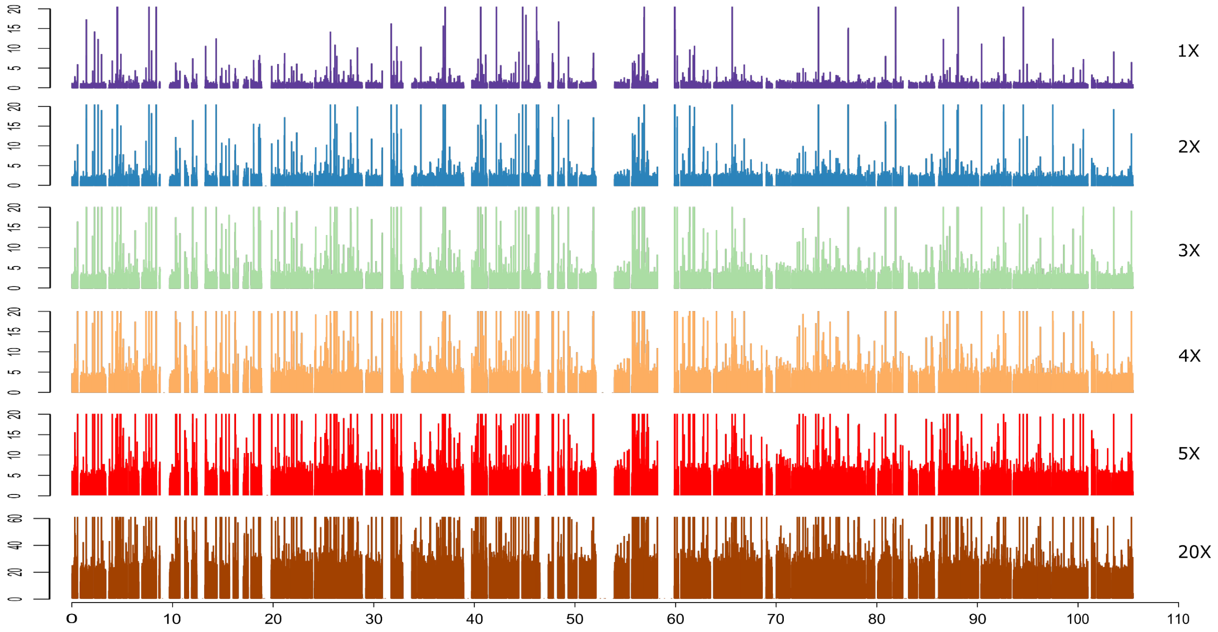


Chromosome 5

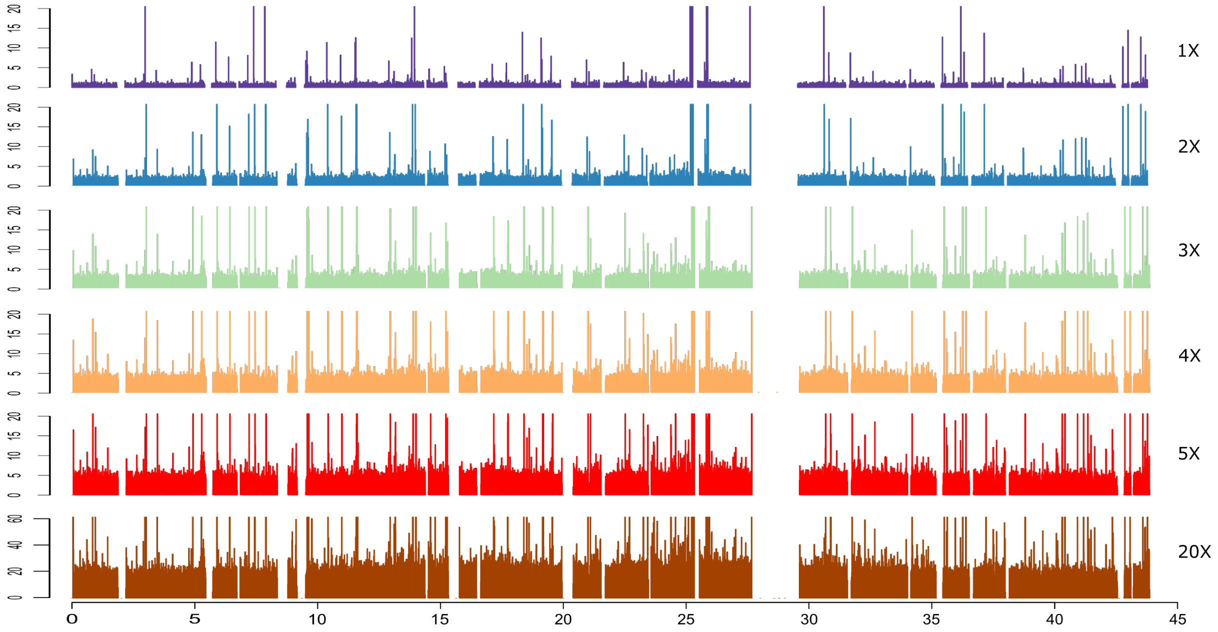

Chromosome 6

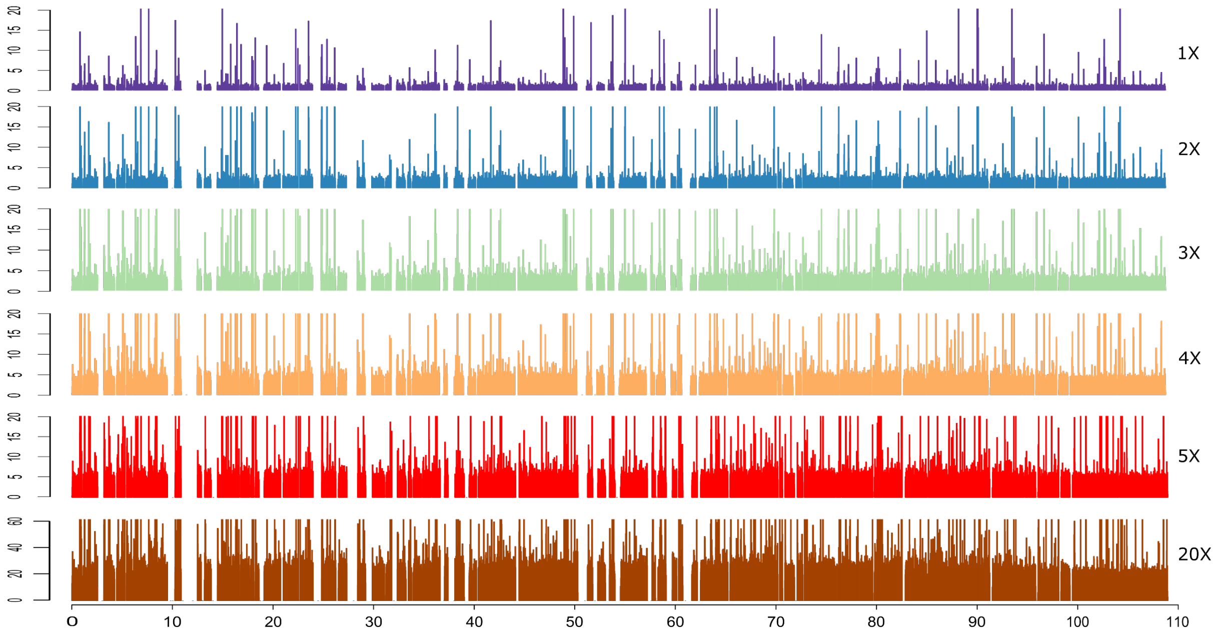


Chromosome 7

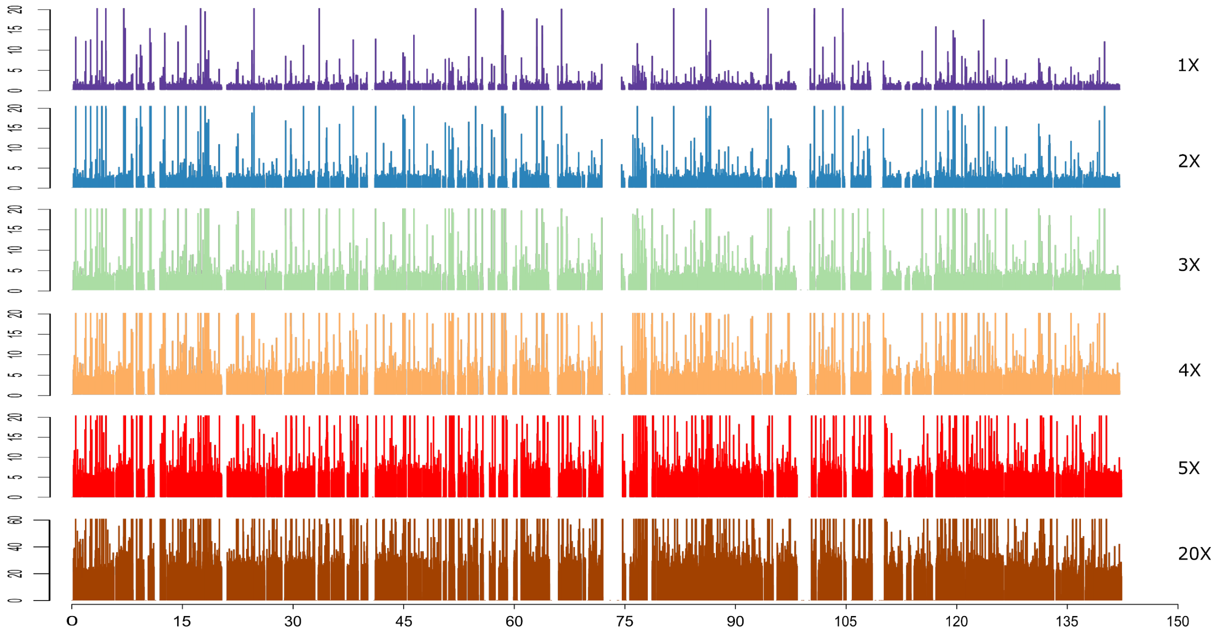

Chromosome 8

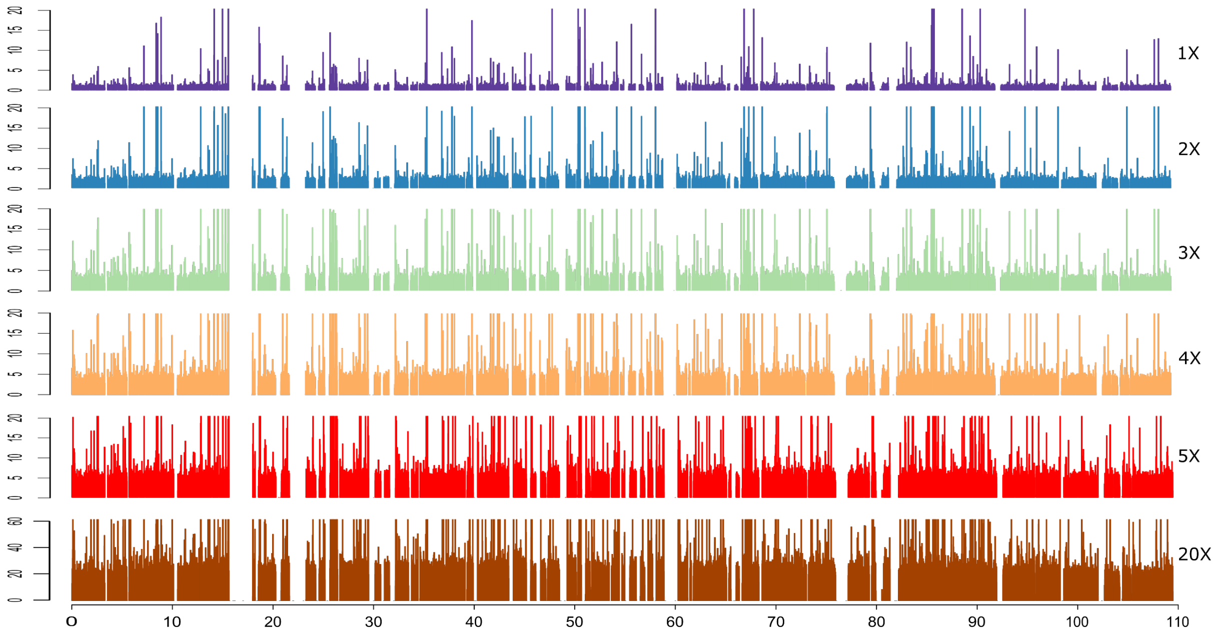


Chromosome 9

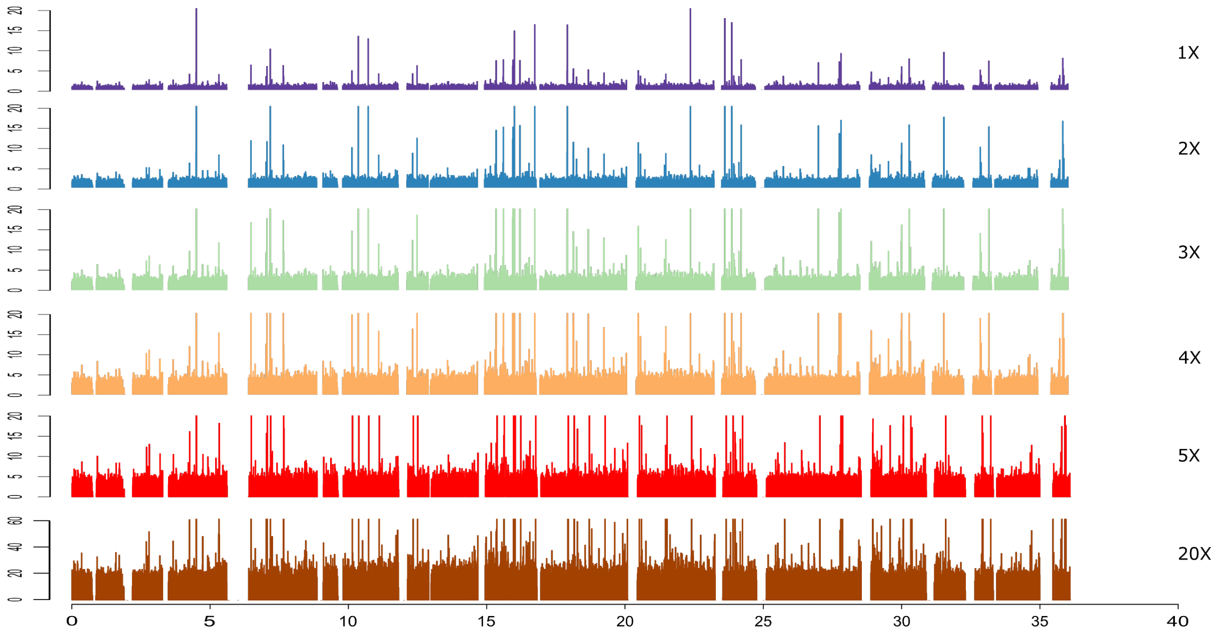

Chromosome 10

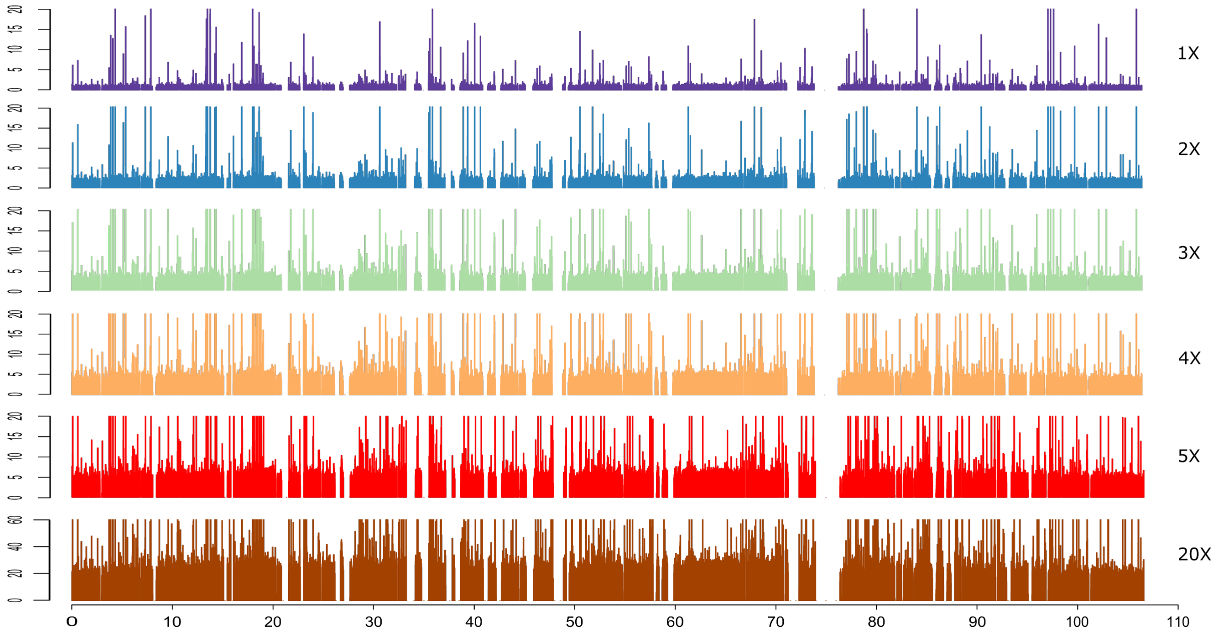


Chromosome 11

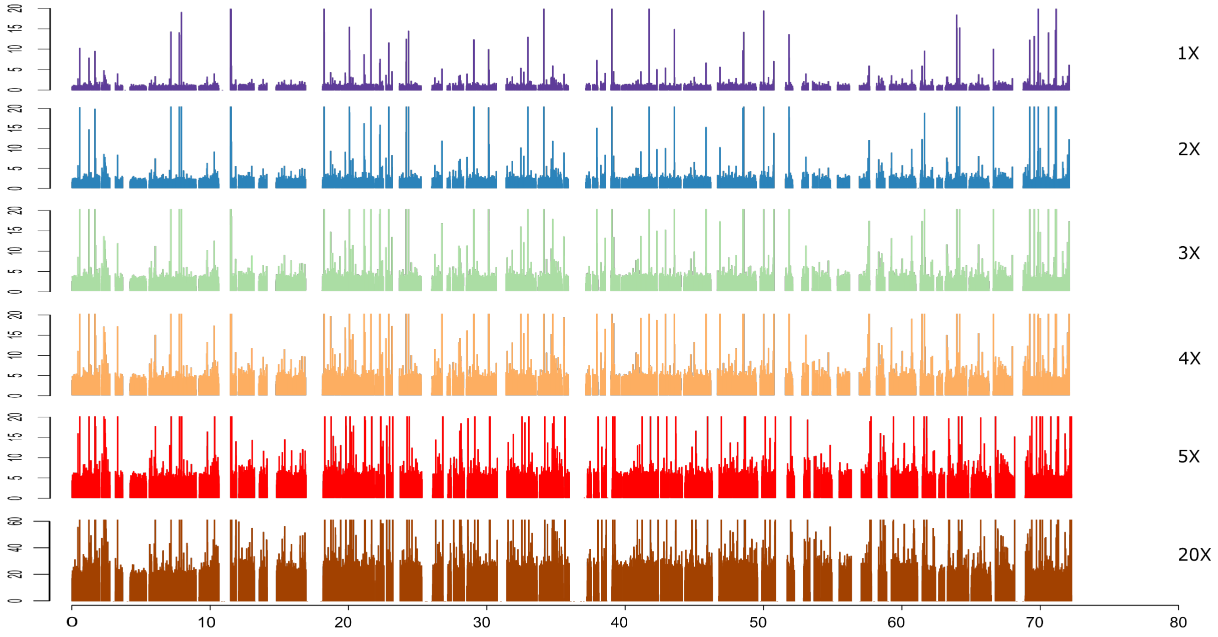

Chromosome 12

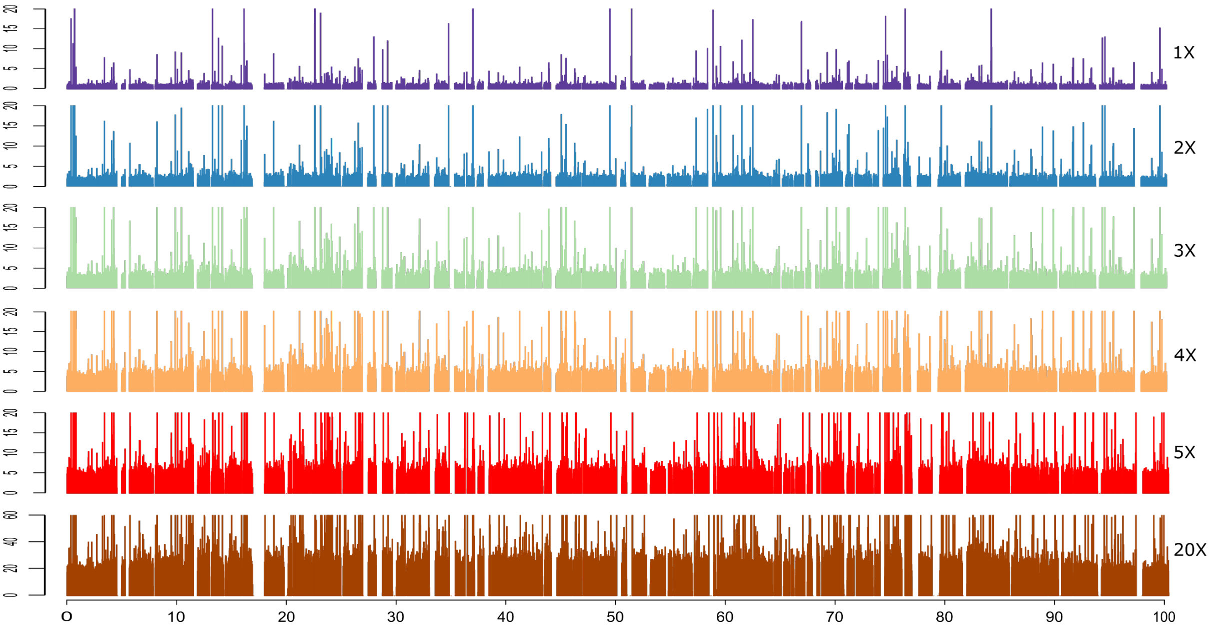
