## Supplementary Data S5 for "Benchmarking of Low Coverage Sequencing Workflows for Precision Genotyping in Eggplant"

**Supplementary Data S5.** Comparative heatmaps depicting the average number of biallelic SNPs identified among the MEGGIC founders across varying sequencing coverages (1X to 5X) and minimum depth of coverage thresholds (from DP 1 to DP 10) using Freebayes and GATK. The colour intensity of the squares reflects the number of polymorphisms identified, with darker shades indicating a higher number of biallelic SNPs.

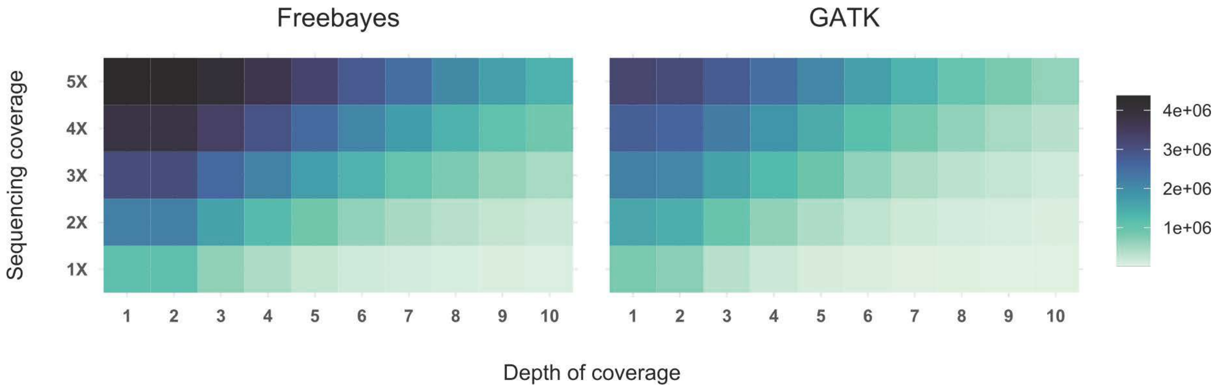
