## Supplementary Data S10 for "Benchmarking of Low Coverage Sequencing Workflows for Precision Genotyping in Eggplant"

**Supplementary Data S10.** Percentage of true positives variants when using different minimum depth of coverage thresholds (from DP 1 to DP 10) across different sequencing coverages (1-5X) using Freebayes. True positive polymorphisms were those shared between the samples and the gold standard. Percentages represent the average across the four S5MEGGIC lines, with SD indicated by error bars (n=20 for 1-4X and n=4 for 5X).

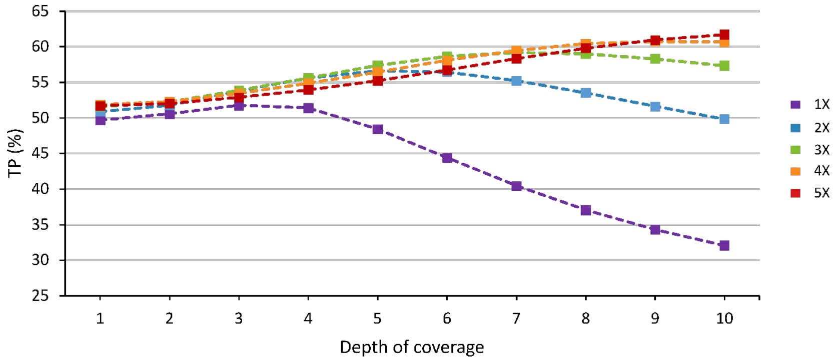
