## Supplementary Data S12 for "Benchmarking of Low Coverage Sequencing Workflows for Precision Genotyping in Eggplant"

**Supplementary Data S12.** Percentage of true positives variants identified in the four S5MEGGIC lines (0% missing data), in three lines (25% missing data), and in two lines (50% missing data). This was evaluated for each combination of minimum depth of coverage thresholds (from DP 1 to DP 10) and sequencing coverages (1-5X).

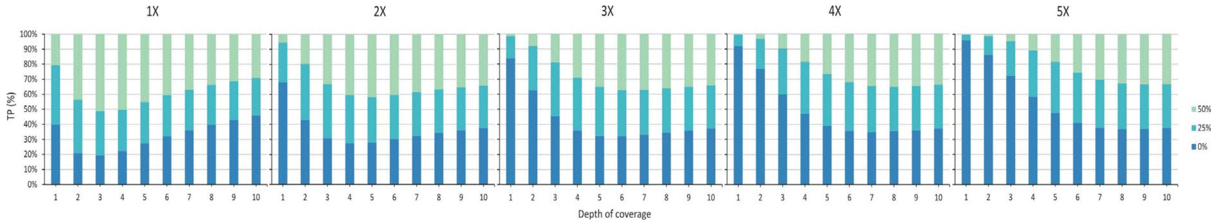
